# HP1 oligomerization compensates for low-affinity H3K9me recognition and provides a tunable mechanism for heterochromatin-specific localization

**DOI:** 10.1101/2021.01.26.428151

**Authors:** Saikat Biswas, Joshua D. Karslake, Ziyuan Chen, Ali Farhat, Peter L. Freddolino, Julie S. Biteen, Kaushik Ragunathan

**Author notes:** equal contribution (Saikat Biswas, Joshua D. Karslake, Ziyuan Chen, Ali Farhat).

## Abstract

HP1 proteins bind with low affinity but high specificity to histone H3 lysine 9 methylation (H3K9me), forming transcriptionally inactive genomic compartments referred to as heterochromatin. How HP1 proteins traverse a complex and crowded chromatin landscape on the millisecond timescale and yet recognize H3K9me with high specificity remains paradoxical. Here, we visualize the single-molecule dynamics of an HP1 homolog, the fission yeast Swi6, in its native chromatin environment. By analyzing the motions of individual Swi6 molecules, we identify mobility states that map to discrete biochemical intermediates. Using mutants that perturb Swi6 H3K9me recognition, oligomerization, or nucleic acid binding, we parse the mechanism by which each biochemical property affects protein dynamics. We find that rather than enhancing chromatin binding, nucleic acid interactions, compete with and titrates Swi6 away from heterochromatin. However, as few as four tandem Swi6 chromodomains are necessary and sufficient to restore H3K9me-dependent localization. Our studies propose propose that HP1 oligomerization stabilizes higher-order protein configurations of a defined stoichiometry that facilitates high-specificity H3K9me recognition and outcompetes the inhibitory effects of nucleic acid-binding.

## INTRODUCTION

Despite having identical genomes, eukaryotic cells can establish distinct phenotypic states that remain stable and heritable throughout their lifetimes (Allis and Jenuwein, 2016). In the context of a multicellular organism, the persistence of epigenetic states is vital to establish and maintain distinct cellular lineages (Bonasio et al., 2010).This process of phenotypic diversification depends, in part, on the post-translational modifications of DNA packaging proteins called histones (Jenuwein and Allis, 2001; Strahl and Allis, 2000). Proteins that can read, write, and erase histone modifications interact weakly and transiently with their histone substrates. Nevertheless, these dynamic, low-affinity interactions between histone modifiers and their substrates encode cellular memories of gene expression that can be inherited following DNA replication and cell division (Stewart-Morgan et al., 2020). H3K9 methylation (H3K9me) is a conserved epigenetic modification associated with transcriptional silencing and heterochromatin formation (Grewal and Moazed, 2003). Heterochromatin establishment is critical for chromosome segregation, sister chromatid cohesion, transposon silencing, and maintaining lineage-specific patterns of gene expression (Nicetto and Zaret, 2019). These diverse cellular functions associated with heterochromatin are critically dependent on the evolutionarily conserved HP1 family of proteins that recognize and bind to H3K9me nucleosomes (Canzio et al., 2014).

HP1 proteins have a distinct architecture that consists of two conserved structural domains: 1) an N-terminal chromodomain (CD) that recognizes H3K9me nucleosomes; and 2) a C-terminal chromoshadow domain (CSD) that promotes oligomerization and mediates heterochromatin-specific protein-protein interactions (**Figure 1A**) (Cowieson et al., 2000). The HP1 CD domain binds to H3K9me peptides with low micromolar affinity (1-10 μM) (Hughes et al., 2007; Jacobs and Khorasanizadeh, 2002). In contrast, the HP1 CSD domain oligomerizes to form higher-order, phase-separated HP1 containing chromatin complexes that exhibit liquid-like properties *in vitro* and *in vivo* (Cowieson et al., 2000; Larson et al., 2017; Sanulli et al., 2018; Strom et al., 2017). A flexible and unstructured hinge region connects the Swi6 CD and CSD domains. The hinge region binds to nucleic acids without any sequence specificity (Smothers and Henikoff, 2001). At present, it is unclear how the competing demands of hinge-mediated nucleic acid binding, CD-dependent H3K9me recognition, and CSD-mediated oligomerization influences HP1 localization *in vivo*.

**Figure 1.**
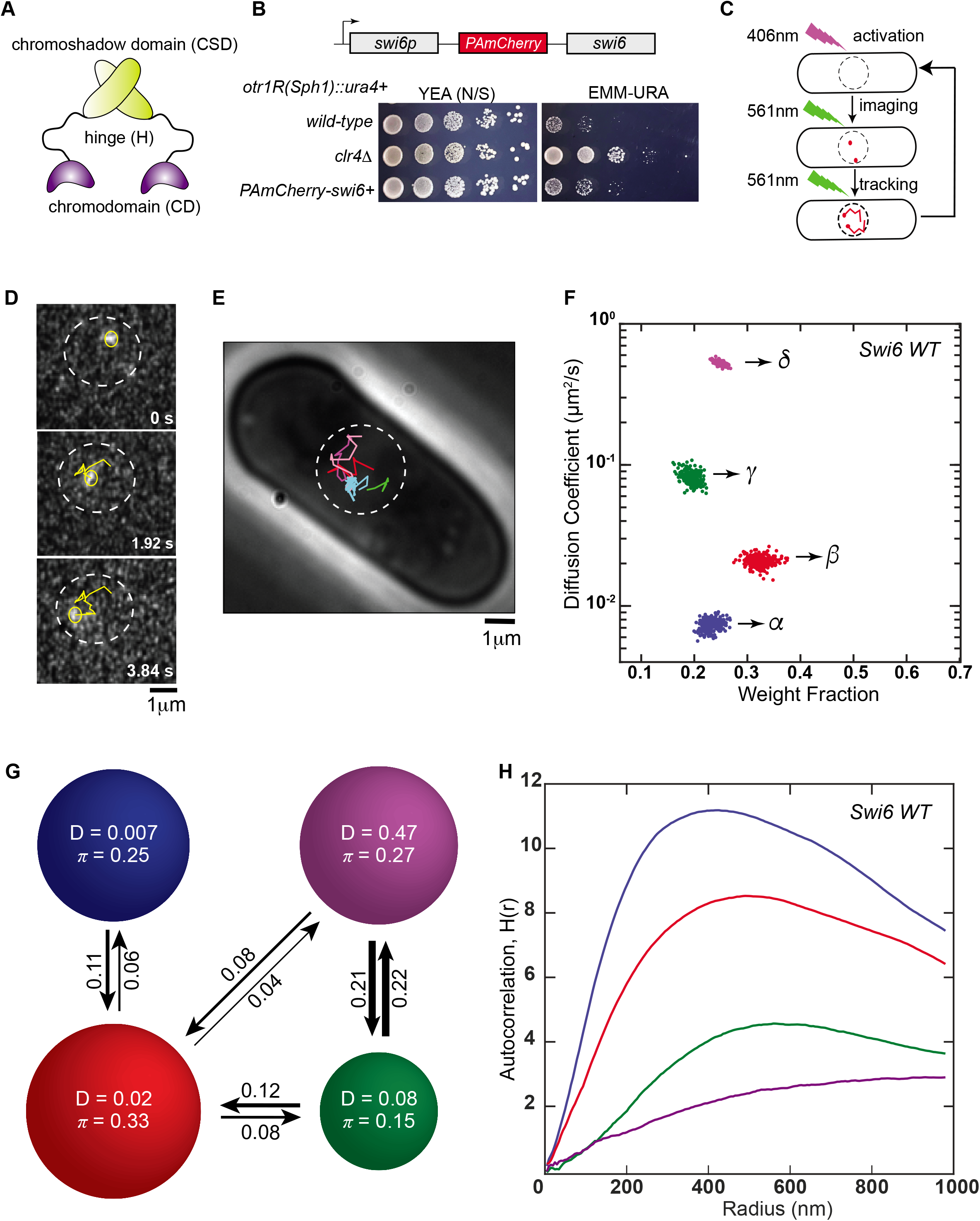
Single particle tracking photoactivation localization microscopy (spt-PALM) of Swi6 in the *S. pombe* nucleus. **1A.** Schematic representation of various domains within Swi6. Each domain has a distinct biochemical property. CD: chromodomain (H3K9me recognition); H: hinge (nucleic acid binding); CSD: chromoshadow domain (protein oligomerization). **1B.** Top: Schematic of the PAmCherry-Swi6 construct showing PAmCherry fused to the N-terminus of Swi6 and expressed from the endogenous *swi6*+ promoter. Bottom: Silencing assay using a *ura4*+ reporter inserted at the pericentromeric repeats (*otr1R*). Cells that establish epigenetic silencing exhibit reduced growth on EMM-URA plates. *clr4Δ* cells lacking H3K9me fail to establish epigenetic silencing and are plated along with PAmCherry-Swi6 expressing cells as a control. **1C.** Single-molecule experiment workflow. After reducing background fluorescence (488 nm, 377 W/cm^2^ for 20 – 40 s), PAmCherry-Swi6 molecules (1 – 3/nucleus) are photoactivated (406 nm, 1.50-4.50 W/cm^2^, 200 ms), then imaged and tracked until photobleaching (561 nm, 71.00 W/cm^2^, 25 frames/sec). The photoactivation/imaging cycle is repeated 10 – 20 times/cell. **1D.** Example trajectory of a single photoactivated PAmCherry-Swi6 molecule. Each frame represents a snapshot of the position of an individual PAmCherry-Swi6 molecule at a different time. Yellow circle: current frame molecule position, yellow line: Swi6-PAmCherry trajectory from photoactivation until the current frame. **1E.** Collection of five example single-particle trajectories in a live *S. pombe* cell. Each color is a single-particle trajectory that is acquired following sequential photoactivation cycles. **1F.** SMAUG identifies four distinct mobility states, (blue *α*, red *β*, green *γ*, and purple *δ*respectively) for PAmCherry-Swi6 in WT cells. Each point represents the average singlemolecule diffusion coefficient vs. weight fraction of PAmCherry-Swi6 molecules in each distinct mobility state at each saved iteration of the Bayesian algorithm after convergence. The dataset contains 10,095 steps from 1491 trajectories. **1G.** Based on the SMAUG identification of four distinct mobility states for PAmCherry-Swi6 in WT cells (four circles with colors as in **Figure 1F** and with average single-molecule diffusion coefficient, *D*, indicated in μm^2^/s), the average probabilities of transitioning between mobility states at each step are indicated as arrows between those two circles, and the circle areas are proportional to the weight fractions, *π*. Low significance transition probabilities below 0.04 are not included. The dataset contains 10,095 steps from 1491 trajectories. **1H.** The Ripley’s analysis exhibits positive autocorrelation values (higher H(r)) for PAmCherry-Swi6 molecules in the slowest (*α*, blue) and the intermediate (*β*, red) state compared to H(*r*) ≤ 2 for molecules in the intermediate (*γ*, green) and fastest (*δ*, purple) state. Each autocorrelation plot is normalized with randomly simulated trajectories from the same state (Methods).

In fission yeast (*Schizosaccharomyces pombe*), H3K9me is enriched at sites of constitutive heterochromatin which includes the pericentromeric repeat sequences (*dg* and *dh*), the telomeres (*tlh*), and the mating-type locus (*mat*) (Allshire and Ekwall, 2015). A conserved SET domain-containing methyltransferase, Clr4, is the sole enzyme that catalyzes H3K9 methylation in *S. pombe* (Ekwall et al., 1996; Ivanova et al., 1998). The major *S. pombe* HP1 homolog, Swi6, senses the resulting epigenetic landscape and binds to H3K9me chromatin with low affinity but high specificity (Ekwall et al., 1995). Swi6 is an archetypal member of the HP1 family of proteins (Canzio et al., 2011). Its ability to simultaneously recognize H3K9me and oligomerize enables Swi6 to spread linearly across broad segments of the chromosome encompassing several hundred kilobases of DNA (Haldar et al., 2011). This process leads to the epigenetic silencing of genes that are distal from heterochromatin nucleation centers.

Based on Fluorescence Recovery After Photobleaching (FRAP) measurements, the turnover rates of Swi6 and other HP1 homologs from sites of heterochromatin range from a few hundred milliseconds to several seconds (Cheutin et al., 2004; Cheutin et al., 2003). This fast turnover could either be due to CD-dependent binding and unbinding of Swi6 from H3K9me nucleosomes, or to CSD-dependent dissociation of Swi6 oligomers. Point mutations that compromise CD binding to H3K9me nucleosomes or CSD mediated oligomerization abolishes Swi6 localization at sites of heterochromatin in *S. pombe* cells (Canzio et al., 2011). Hence, there is a need to quantitatively delineate the respective contributions of oligomerization and H3K9me recognition towards Swi6 localization.

At equilibrium, the fraction of Swi6 molecules bound to constitutive heterochromatin in *S. pombe* cells comprises only about a very small fraction of the total pool (Cheutin et al., 2004). Hence, the vast majority of Swi6 molecules are located elsewhere in the genome potentially engaged in promiscuous DNA- and chromatin-dependent interactions (Keller et al., 2012; Stunnenberg et al., 2015). How Swi6 molecules are partitioned between different bound intermediates, and how these relative occupancy changes affect Swi6 localization at sites of H3K9me, remain unclear. Indeed, Swi6 overexpression enhances epigenetic silencing of a reporter gene, and this silent state is heritable during mitosis and meiosis (Nakayama et al., 2000). Therefore, Swi6 is a dose-sensitive heterochromatin associated factor, and mechanisms that alter its occupancy at sites of H3K9me can have a profound impact on transcriptional silencing and epigenetic inheritance. *In vitro* experiments cannot reproduce the intricate balance of competing and complementary binding interactions in cells that give rise to such complex outcomes.

*In vitro* binding measurements have thus far served as the gold standard to measure interactions between chromatin readers and modified histone peptides or recombinant nucleosome substrates (Canzio et al., 2011; Canzio et al., 2013; Hiragami-Hamada et al., 2016; Isaac et al., 2017; Kilic et al., 2015; Nishibuchi et al., 2014; Sanulli et al., 2019). These studies are typically carried out under dilute, non-competitive solution conditions, which do not reflect the chromatin environment that Swi6 encounters in the nucleus. For example, in fission yeast cells, only ~2% of the total number of nucleosomes in the genome are marked with H3K9me (http://www.pombase.org). Therefore, the vast excesses of DNA and nucleosomes in the fission yeast genome have no H3K9 methylation (H3K9me0) and thus function as competitive inhibitors of Swi6 target search. In this study, we use single-particle tracking photoactivated localization microscopy (spt-PALM) to measure the *in vivo* binding dynamics of Swi6 in real-time as it samples the fission yeast nucleus (Manley et al., 2008; Tuson and Biteen, 2015). By analyzing the trajectories of individual Swi6 molecules with high spatial and temporal resolution, we delineate the biochemical attributes of Swi6 that give rise to distinct mobility states. These measurements enabled us to engineer precise degrees of multivalency within Swi6 that entirely circumvent the need for CSD-dependent oligomerization while suppressing promiscuous nucleic acid binding. We find that the simultaneous engagement of at least four H3K9me CD domains is both necessary and sufficient for the heterochromatin-specific targeting of Swi6. Overall, our results demonstrate that the evolutionarily conserved process of HP1 oligomerization may represent a tunable mechanism that compensates for weak, low-affinity H3K9me recognition while still enabling the formation of stable and heritable epigenetic states.

## RESULTS

### Single-molecule dynamics of Swi6 in live *S. pombe* indicate a heterogeneous environment

Because Swi6 molecules within the fission yeast nucleus undergo binding and unbinding events in a complex environment, we used single-molecule tracking to measure the resulting heterogeneous dynamics. For instance, a Swi6 molecule bound to an H3K9me nucleosome is on average likely to exhibit slower, more confined motion compared to rapidly diffusing proteins. Hence, the biochemical properties of Swi6 will directly influence its mobility within the fission yeast nucleus. We transformed *S. pombe* cells with PAmCherry fused to the N-terminus of Swi6 (Subach et al., 2009). This fusion protein replaces the wild-type endogenous Swi6 gene and serves as the sole source of Swi6 protein in fission yeast cells (**Figure 1B**). To test whether PAmCherry-Swi6 is functional to establish epigenetic silencing, we used strains where a *ura4*+ reporter is inserted within the outermost pericentromeric repeats (*otr1R*) (Ekwall et al., 1997). The *ura4*+ gene is required for uracil biosynthesis. If *ura4*+ is silenced, cells exhibit reduced growth on medium lacking uracil (-URA) (**Figure 1B**). Like wild-type Swi6 expressing cells, cells expressing PAmCherry-Swi6 exhibit reduced growth on –URA medium (**Figure 1B**). As a control, we plated cells lacking the H3K9 methyltransferase Clr4 (*clr4*Δ). These cells exhibit prolific growth consistent with *ura4*+ being expressed due to the loss of H3K9me dependent epigenetic silencing (**Figure 1B**). Therefore, we concluded that the PAmCherry-Swi6 resembles the wild-type Swi6 protein in its ability to establish epigenetic silencing.

We used single-molecule microscopy to investigate the dynamics of Swi6 molecules with high spatial (10 – 20 nm) and temporal (40 ms) resolution (Tuson and Biteen, 2015). We photoactivated one PAmCherry-Swi6 fusion protein per nucleus with 406-nm light. Next, we imaged the photoactivated Swi6 molecules with 561-nm laser excitation until photobleaching, obtain a 5-15 step trajectory based on localizing molecules at 40 ms intervals, then repeated the photoactivation/imaging cycle with another PAmCherry-Swi6 molecule ~10 times per cell (**Figure 1C-D**). We observed both stationary and fast-moving molecules (**Figure 1E**). We hypothesized that each type of motion, which we term a “*mobility state*”, corresponds to a distinct biochemical property of Swi6 in the cell (e.g., bound vs unbound) and that molecules can transition between the different mobility states during a single trajectory. Thus, rather than assign a single diffusion coefficient to each single-molecule trajectory, we performed a single-step analysis with the Single-Molecule Analysis by Unsupervised Gibbs sampling (SMAUG) algorithm (**Methods**) (Karslake et al., 2020). SMAUG estimates the biophysical descriptors of a system by embedding a Gibbs sampler in a Markov Chain Monte Carlo framework. This nonparametric Bayesian analysis approach determines the most likely number of mobility states and the average diffusion coefficient of single molecules in each state, the occupancy of each state, and the probability of transitioning between different mobility states between subsequent 40 ms frames.

For each cell, we sampled over a thousand trajectories of PAmCherry-Swi6 within the nucleus of *S. pombe* cells (**Supplementary Figure 1A**). SMAUG analyzed the ~10,000 steps from these trajectories in aggregate. For cells expressing fusions of PAmCherry-Swi6, the algorithm converged to four mobility states and estimated the diffusion coefficient, *D*, and the fraction of molecules in each state, *π*, for each iteration (each dot in Figure 1F is the assignment from one iteration of the SMAUG analysis). The dynamics of Swi6 in the nucleus are best described by four mobility states, and each mobility state has a different average diffusion coefficient, *D*_avg_ (we refer to the states as *α, β, γ*, and *δ* in order of increasing *D*_avg_ in **Figure 1F**). The clusters of points correspond to estimates of D and π for each of the four mobility states and the spread in the clusters indicates the inferential uncertainty. The slowest mobility state comprises ~25% of the Swi6 molecules; its average diffusion coefficient (*D*_avg,*α*_ = 0.007 ± 0.001 μm^2^/s) is close to the localization precision of the microscope, indicating no measurable motion for these molecules (error bars indicate the 95% credible interval). Given that Swi6 forms discrete foci at sites of constitutive heterochromatin (centromeres, telomeres and the mating-type locus), the *α* mobility state likely corresponds to Swi6 molecules that are stably bound at sites of H3K9me.

On the other hand, the fastest diffusion coefficient we measured (*D*_avg, *δ*_ = 0.51 ±0.02 μm^2^/s) matches that of proteins that freely diffuse within the nucleus (Rappoport et al., 2009). Therefore, the fastest mobility *δ* state likely describes unbound Swi6 molecules that do not interact with chromatin. We also measured how often Swi6 molecules in one mobility state transition to another mobility state. We discovered that Swi6 molecules are more likely to transition between adjacent rather than non-adjacent mobility states, demonstrating that a distinct hierarchy of biochemical interactions dictates how Swi6 locates sites of H3K9me in the fission yeast genome (**Figure 1G**). Notably, only molecules in the *β* intermediate mobility state transition with high probability to the slow mobility *α* state, which (as noted above) we assign to the Swi6 molecules stably bound at sites of H3K9me.

We used a spatial auto-correlation analysis to measure inhomogeneities in Swi6 mobility patterns (**Figure 1H**). The Ripley’s H-function, H(*r*), measures deviations from spatial homogeneity for a set of points and quantifies the correlation as a function of the search radius, *r* (Ripley, 1976). We calculated H(*r*) values for the positions of molecules in each mobility state from the single-molecule tracking dataset. To eliminate the bias that would come from the spatial correlation between steps along the same trajectory, we compared the experimental observations to a null model by randomly simulating confined diffusion trajectories for each of the four mobility states (**Methods**). We represented the spatial autocorrelation values associated with each state with an H(*r*) function normalized using simulated trajectories (**Supplementary Figure 1B-C**). The real trajectories show both significantly higher magnitude and longer distance correlations than a realistic simulated dataset, demonstrating that Swi6 mobility is indeed heterogeneous in the nucleus.

We observed low autocorrelation values for the *δ* and γ states (purple and green) and high autocorrelation values for the *β* and *α* states (blue and red) (**Figure 1H**). The crosscorrelation between the *β, γ*, and *δ* mobility states and the slow-moving *α* state further supports our model of spatial confinement: molecules in the *β* state are most likely to be proximal to H3K9me-bound molecules in the *α* state (**Supplementary Figure 1D**). In summary, the combination of transition plots and spatial homogeneity maps suggests that the *β* and *γ* mobility states represent biochemical intermediates that sequester Swi6 molecules from transitioning between the fully bound *α* state and the fast diffusing, chromatin-unbound *δ* state.

### The slowest Swi6 mobility state corresponds to its localization at sites of heterochromatin formation

Following the baseline characterization of the different mobility states associated with Swi6 and their relative patterns of spatial confinement, we used fission yeast mutants to dissect how each biochemical property of Swi6 gives rise to a unique mobility state. As a first approximation, we hypothesized that the major features likely to affect Swi6 binding within the nucleus are: 1) CD-dependent H3K9 methylation recognition, 2) hinge-mediated nucleic acid binding, and 3) CSD-mediated oligomerization (Canzio et al., 2014).

We deleted the sole *S. pombe* H3K9 methyltransferase, Clr4, to interrogate how the four mobility states associated with Swi6 diffusion respond to the genome-wide loss of H3K9 methylation. The mobility of PAmCherry-Swi6 is substantially different in (H3K9me0) *clr4Δ* cells compared to *clr4+* cells. Most prominently, the majority of Swi6 molecules in *clr4Δ* cells move rapidly and show no subnuclear patterns of spatial confinement compared to wild-type cells. The slow mobility *α* state is absent, and the fraction of molecules in the intermediate mobility *β*state decreases two-fold (**Figure 2A**). Hence, both the *α* and *β* states depend on Swi6 binding to H3K9me nucleosomes, with *α* occurring exclusively in *clr4+* cells and *β* apparently representing a mixture of H3K9me-dependent and H3K9me-independent substates (present in *clr4+* and *clr4Δ* cells). In contrast, the weight fractions of the *γ* and *δ* states substantially increase in H3K9me0 *clr4Δ* cells suggesting that these mobility states are exclusively H3K9me-independent (**Figure 2A**).

**Figure 2.**
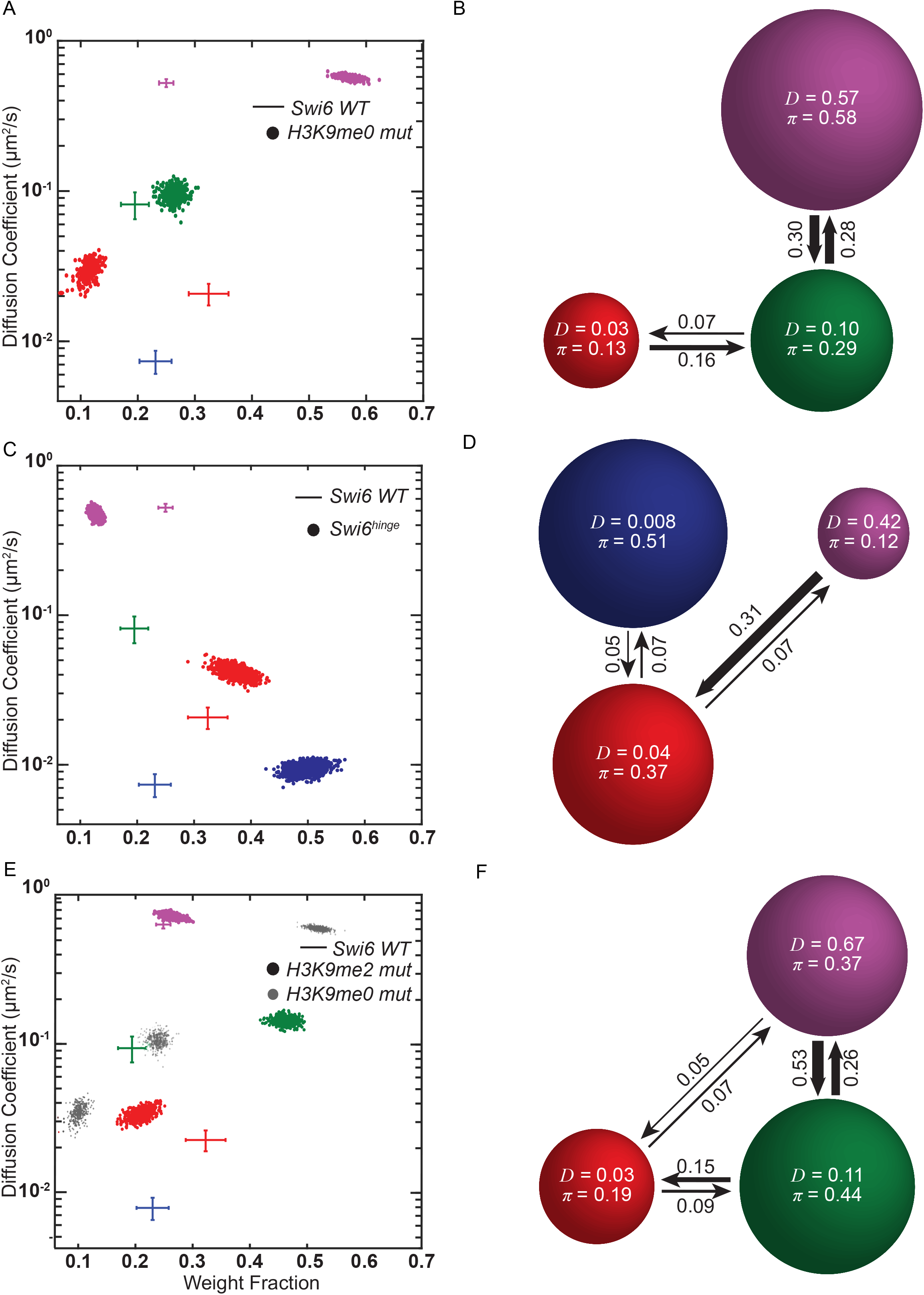
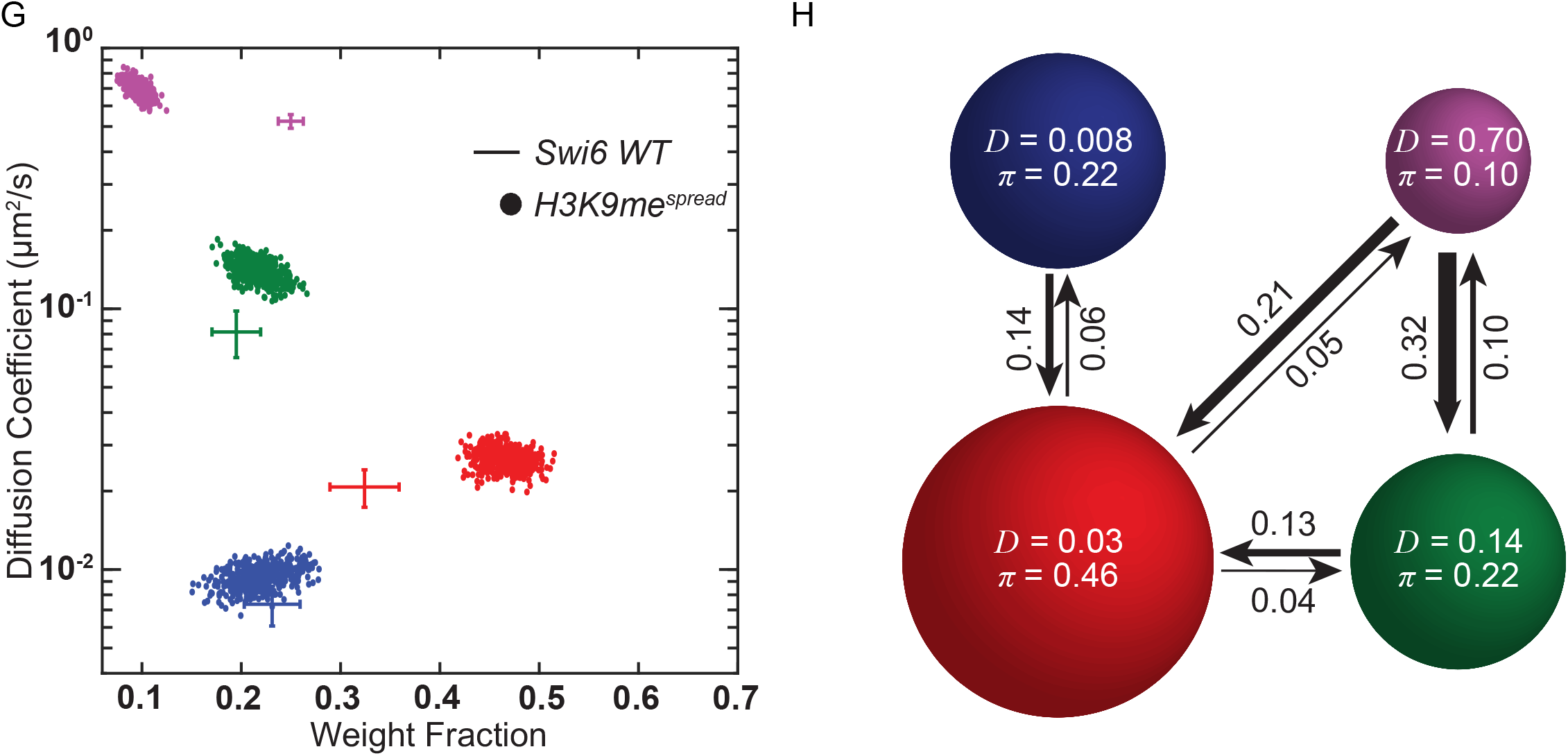
Individual Swi6 mobility states reflect distinct biochemical intermediates. **2A.** Average single-molecule diffusion coefficients and weight fraction estimates for PAmCherry-Swi6 molecules expressed in H3K9me0 mut (*clr4*Δ) cells. SMAUG identifies three distinct mobility states, (red *β*, green *γ*, and purple *δ*, respectively) for PAmCherry-Swi6 in H3K9me0 mut cells. Each point represents the average single-molecule diffusion coefficient vs. weight fraction of PAmCherry-Swi6 molecules in each distinct mobility state at each saved iteration of the Bayesian algorithm after convergence. The dataset contains 10432 steps from 2463 trajectories. Specifically, the *α* mobility state is absent in H3K9me0 mut (*clr4*Δ) cells. The average and standard deviation of the wild-type Swi6 clusters (**Figure 1F**) are provided as a reference (cross hairs). **2B.** Based on the SMAUG identification of three distinct mobility states for PAmCherry-Swi6 in H3K9me0 mut cells (three circles with colors as in **Figure 2A** and with average single-molecule diffusion coefficient, D, indicated in μm^2^/s), the average probabilities of transitioning between mobility states at each step indicated as arrows between those two circles, and the circle areas are proportional to the weight fractions, *π*. Low significance transition probabilities below 0.04 are not included. The dataset contains 10432 steps from 2463 trajectories. **2C.** Average single-molecule diffusion coefficients and weight fraction estimates for PAmCherry-Swi6^hinge^ expressing cells (*swi6 KR25A* mutant, in which 25 lysine and arginine residues in the Swi6 hinge region are replaced with alanine). SMAUG identifies three distinct mobility states, (blue *α*, red *β* and purple *δ*, respectively) for PAmCherry-Swi6^hinge^ cells Each point represents the average single-molecule diffusion coefficient vs. weight fraction of PAmCherry-Swi6 molecules in each distinct mobility state at each saved iteration of the Bayesian algorithm after convergence. The dataset contains 12788 steps from 1210 trajectories. Specifically, the *γ* mobility state is absent in PAmCherry-Swi6^hinge^ expressing cells. The wild-type Swi6 clusters (**Figure 1F**) are provided for reference (cross hairs). **2D.** Based on the SMAUG identification of three distinct mobility states for PAmCherry-Swi6 in Swi6^hinge^ cells (three circles with colors as in **Figure 2C** and with average single-molecule diffusion coefficient, D, indicated in μm^2^/s), the average probabilities of transitioning between mobility states at each step are indicated as arrows between those two circles, and the circle areas are proportional to the weight fractions, *π*. Low significance transition probabilities below 0.04 are not included. The dataset contains 12788 steps from 1210 trajectories. **2E.** Average single-molecule diffusion coefficients and weight fraction estimates for PAmCherry-Swi6 molecules expressed in H3K9me2 mut cells (*clr4 F449Y* mutant which has a phenylalanine to tyrosine substitution within the Clr4 catalytic SET domain). SMAUG identifies three distinct mobility states, (red *β*, green *γ*, and purple *δ*, respectively) for H3K9me2 mut cells. Each point represents the average single-molecule diffusion coefficient vs. weight fraction of PAmCherry-Swi6 molecules in each distinct mobility state at each saved iteration of the Bayesianalgorithm after convergence. The dataset contains 14837 steps from 2308 trajectories. The wild-type Swi6 clusters (**Figure 1F**) and H3K9me0 mut (*clr4*Δ) (**Figure 2A**) are provided as a reference cross hairs and grey circles, respectively. **2F.** Based on the SMAUG identification of three distinct mobility states for PAmCherry-Swi6 in H3K9me2 mut cells (three circles with colors as in **Figure 2E** and with average single-molecule diffusion coefficient, D, indicated in μm^2^/s), the average probabilities of transitioning between mobility states at each step are indicated as arrows between those two circles, and the circle areas are proportional to the weight fractions, *π*. Low significance transition probabilities below 0.04 are not included. The dataset contains 14837 steps from 2308 trajectories. **2G.** Average single-molecule diffusion coefficients and weight fraction estimates for PAmCherry-Swi6 molecules in H3K9me^*spread*^(*mst2Δ, epe1Δ*) cells. SMAUG identifies four distinct mobility states, (blue *α*, red *β*, green *γ*, and purple *δ*, respectively) for PAmCherry-Swi6 in H3K9me^*spread*^ cells. Each point represents the average single-molecule diffusion coefficient vs. weight fraction of PAmCherry-Swi6 molecules in each distinct mobility state at each saved iteration of the Bayesian algorithm after convergence. The dataset contains 11425 steps from 1287 trajectories. The wild-type Swi6 diffusion coefficients (**Figure 1F**) are provided as a reference (cross hairs). **2H.** Based on the SMAUG identification of four distinct mobility states for PAmCherry-Swi6 in H3K9me^*spread*^ cells (four circles with colors as in **Figure 2G** and with average single-molecule diffusion coefficient, *D*, indicated in μm^2^/s), the average probabilities of transitioning between mobility states at each step are indicated as arrows between those two circles, and the circle areas are proportional to the weight fractions. Low significance transition probabilities below 0.04 are not included. The dataset contains 11425 steps from 1287 trajectories.

A tryptophan to alanine substitution (W104A) within the Swi6 chromodomain (CD) attenuates H3K9me binding approximately 100-fold (Swi6 CD^mut^) (Canzio et al., 2013; Jacobs and Khorasanizadeh, 2002). We expressed PAmCherry-Swi6 CD^mut^ in *S. pombe* cells. Our SMAUG analysis identified three mobility states for PAmCherry-Swi6 CD^mut^ (**Supplementary Figure 2A**), and the distribution of mobility states is similar to that of Swi6 in H3K9me0 *clr4Δ* cells (c.f. **Figure 2A**). Furthermore, the spatial autocorrelation of the different mobility states validates the loss of confinement of Swi6 molecules at sites of constitutive heterochromatin in *clr4Δ* cells. The majority of Swi6 molecules in H3K9me0 *clr4Δ* cells exhibited low H(*r*) values, indicating a lack of spatial correlation in the absence of H3K9me (**Supplementary Figure 2B**) as opposed to the spatially correlated motions we observe in the case of wild-type Swi6 (**Figure 1H**). We also estimated transition probabilities for the different mobility states based on our experimental data. In the absence of the H3K9me dependent low-mobility *α* state, PAmCherry-Swi6 molecules predominantly reside in and transition between the *γ* and *δ* fast-mobility states with only rare transitions to the *β* state (**Figure 2B**).

### Nucleic acid binding defines an intermediate Swi6 mobility state

All HP1 proteins have a variable-length hinge region that connects the H3K9me recognition CD domain and the CSD oligomerization domain (Bryan et al., 2017). In the case of Swi6, the hinge region has twenty-five basic lysine and arginine residues that modulate the interaction between Swi6 and nucleic acids (DNA and RNA). We replaced all twenty-five lysine and arginine residues with alanine (Swi6^hinge^) and imaged the mobility patterns of PAmCherry-Swi6^hinge^ (Keller et al., 2012). Neutralizing the net positive charge within the hinge region results in fewer fast diffusing molecules and a substantially larger proportion of spatially restricted Swi6^hinge^ molecules relative to the wild-type protein. These qualitative observations were consistent with our quantitative analysis by SMAUG. We detect three mobility states (as opposed to four in the case of the wild-type Swi6 protein), a substantial increase in the populations of the H3K9me dependent *α* slow mobility and *β* intermediate mobility states and a concomitant decrease in the population of the *δ* fast diffusing state (**Figure 2C**). Notably, the *γ* intermediate mobility state is absent in the case of PAmCherry-Swi6^hinge^ (**Figure 2C**). Hence, we conclude that the *γ* intermediate mobility state corresponds to Swi6 nonspecifically bound to nucleic acids (DNA or RNA). The two-fold increase in the weight fraction of the slow *α* state suggests that, in the absence of DNA or RNA binding, PAmCherry-Swi6^hinge^ molecules preferentially remain bound to H3K9me chromatin. In addition, the diffusion coefficient of the *β* mobility state exhibits an increase suggesting that nucleic acid binding stabilizes this intermediate.

Based on the assignment of transition probabilities between the mobility states, we identified two critical features of Swi6 dynamics in the hinge mutant: 1) PAmCherry-Swi6^hinge^ molecules in the slow mobility *α* state transition less often to the remaining *β* and *δ* mobility states, and 2) the probability of cross-transitions between the fast diffusing *δ* state and the intermediate mobility *β* state increases (**Figure 2D**). These transitions, although negligible in the case of the wild-type PAmCherry-Swi6 protein, become prominent in the case of PAmCherry-Swi6^hinge^ mutant. By eliminating nucleic acid binding, we observed a substantial increase in H3K9me dependent and H3K9me independent chromatin association. Overall, these measurements strongly suggest that nucleic acid binding interactions drive Swi6 dissociation from chromatin and compete with CD-dependent H3K9me recognition.

### Weak H3K9me nucleosome interactions result in an intermediate Swi6 mobility state

Deleting Clr4 reduces but does not eliminate the population of the *β* intermediate mobility state (**Figure 2A**). Therefore, this *β* state combines both H3K9me-dependent and H3K9me--independent components. We hypothesized that the *β* intermediate mobility state likely represents a transient sampling of chromatin by Swi6 (H3K9me or H3K9me0) before its stable binding at sites of H3K9me (*α* state). Swi6 binds to H3K9me3 chromatin with higher affinity compared to other H3K9me states (H3K9me1/2). To eliminate the high affinity Swi6 binding state and exclusively interrogate transient chromatin interactions, we replaced the H3K9 methyltransferase, Clr4, with a mutant methyltransferase (Clr4 F449Y, referred to here as the Clr4 H3K9me2 mutant) that catalyzes H3K9 mono- and di-methylation (H3K9me1/2) but is unable to catalyze tri-methylation (H3K9me3) due to a mutation within the catalytic SET domain. We verified the expression of the Clr4 mutant protein in fission yeast cells using a *myc* epitope tag. Following single particle tracking measurements of Swi6, we found that cells expressing the Clr4 H3K9me2 mutant exhibit only three mobility states (**Figure 2E**), having lost the *α* state. These results are consistent with our expectations and previously published data where selectively eliminating H3K9me3 interrupts stable Swi6 association at sites of heterochromatin formation (Jih et al., 2017). We also observed a two-fold increase in the weight fraction of molecules residing in the *β* intermediate mobility state relative to H3K9me0 cells, suggesting that the presence of H3K9me2 is sufficient to increase the occupancy of the Swi6 chromatin-bound state (**Figure 2E**). Mapping the transition probabilities of Swi6 molecules between the remaining three mobility states further supports our conclusions. We observed a substantial increase in the transition probability between the *δ* and *γ* mobility states, which is negligible or absent in H3K9me0 cells (**Figure 2F**). Therefore, H3K9me2 enhances Swi6 chromatin binding but is incapable of driving Swi6 occupancy to the immobile *α* state.

Simultaneously, we also tested how H3K9me spreading mutants, which enhance heterochromatin associated silencing, affect Swi6 dynamics. We focused on two negative regulators: 1) Epe1, a putative H3K9 demethylase that erases H3K9me, and 2) Mst2, an H3K14 acetyltransferase that acetylates histones and promotes active transcription (Ayoub et al., 2003; Reddy et al., 2011; Zofall and Grewal, 2006). We deleted either *epe1* or *mst2* individually in cells expressing PAmCherry-Swi6 (**Supplementary Figure 2C-D**). We observed relatively few changes in the weight fractions of the different Swi6 mobility states in these individual mutants. However, simultaneously deleting both *epe1* and *mst2* leads to a more dramatic rearrangement of the Swi6 mobility states (**Figure 2G**). We refer to this double mutant as the H3K9me^spreading^ mutant. The fraction of molecules in the *β* intermediate mobility state increases nearly two-fold in the H3K9me^spreading^ mutant, which also coincides with a near-complete depletion of Swi6 molecules from the unbound *δ* state (**Figure 2G**).

We measured transition probabilities between the different mobility states in the H3K9me^spreading^ mutant cells (**Figure 2H**). We observed a significant increase in cross-transitions between the fast diffusing *δ* state and the intermediate *β* mobility state, suggesting that the reorganization of the H3K9me epigenome enhances chromatin binding and makes direct binding of Swi6 to histones more likely without the need for a DNA-binding intermediate. Under our imaging conditions, we detected a negligible number of direct transitions from the fast mobility *δ* state to the slow mobility *α* state. As expected, deleting *clr4* in this H3K9me^spreading^ mutant (H3K9me^spreading^ *clr4Δ*) collapses the *β* intermediate mobility state from 48% to ~14%, similar to what we observed in *epe1*+ *mst2*+ *clr4Δ* cells (**Supplementary Figure 2E and Figure 2A**).

By synthesizing our results using the H3K9me2 mutant and the H3K9me^spreading^ mutant strains, we conclude that the *β* intermediate corresponds to a chromatin sampling state consisting of weak and unstable interactions between Swi6 and H3K9me or H3K9me0 nucleosomes. These transient chromatin interactions increase in the case of H3K9me^spreading^ mutants and decrease in the case of Clr4 H3K9me2 mutants, and the simultaneous presence of both mutations shows the latter phenotype.

### Simulating Swi6 mobility state transitions reveals H3K9me binding specificity *in vivo*

Our fission yeast mutants enabled us to assign biochemical properties to each of our experimentally determined mobility states (**Figure 3A**). The state-to-state transitions inferred by SMAUG (**Figure 1F**) estimate the probability with which a molecule of Swi6 assigned to one state during one 40-ms imaging frame will be assigned to some other state during the subsequent imaging frame. However, the SMAUG inferences are limited by the timescale of the experiment (40 ms/imaging frame, which is selected to optimize signal-to-noise and track length due to photobleaching). Thus, transitions between mobility states that occur faster than the time resolution of the experiment may not be detected (e.g., rapid transitions from *α➔β* and then *β➔γ* may be misclassified as a single *α➔γ* transition). To infer the chemical rate constants underlying the observed state transitions of Swi6, we implemented a high temporal resolution model that uses approximate Bayesian computation (ABC) to determine the set of fine-scale chemical rate constants most consistent with the transition matrices obtained experimentally via SMAUG (Beaumont et al., 2002; Csillery et al., 2010).

**Figure 3.**
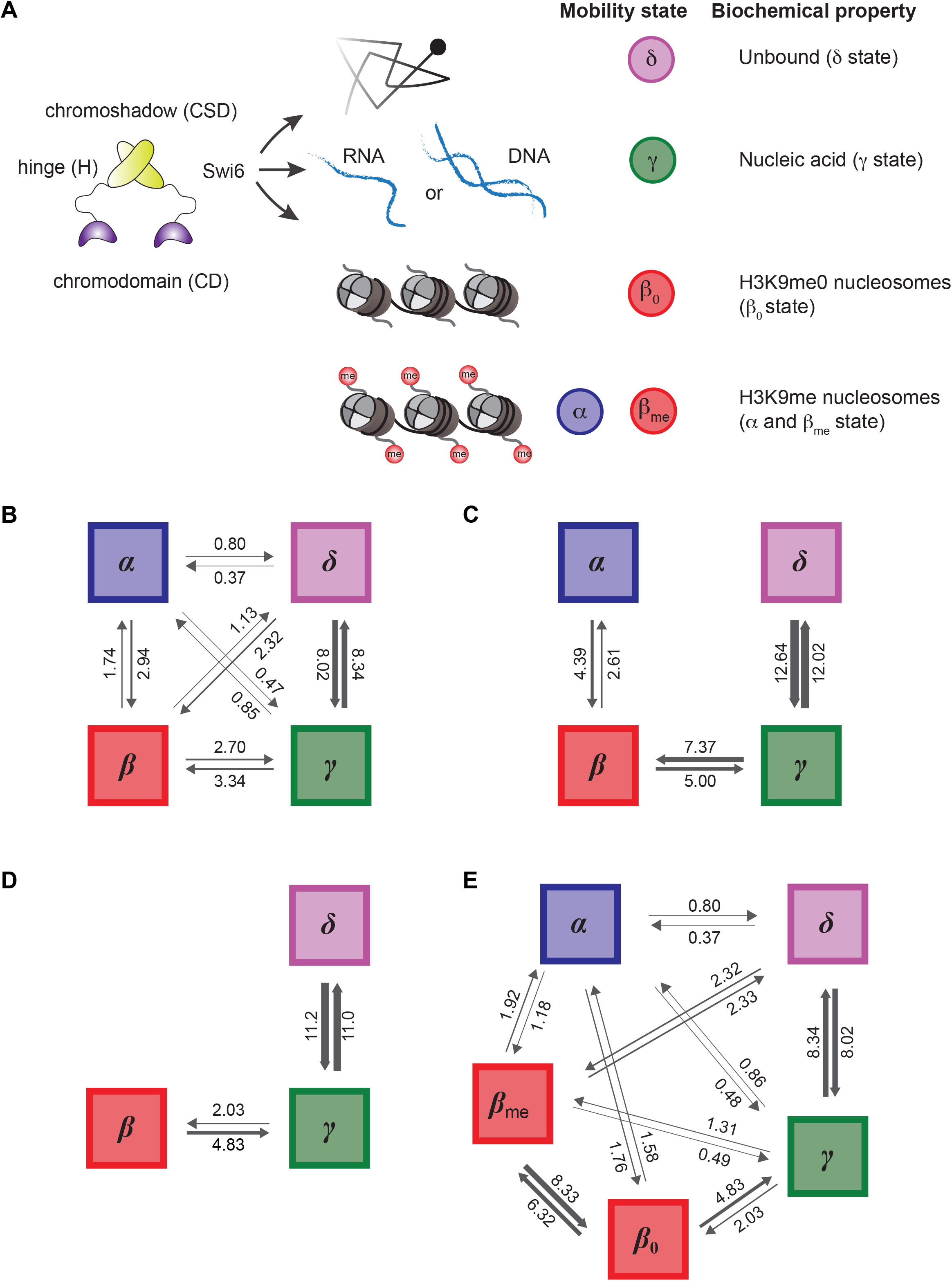
Rate constants inferred from fine-grained chemical kinetic simulation and approximate Bayesian computation. **3A.** Schematic showing the inferred biochemical nature of each state considered in our finegrained chemical simulations. **3B.** Inferred rate constants for cells expressing wild type Swi6, in units of 1/sec, for transitions of Swi6 assuming that there are four biochemical states of Swi6 and that any state can chemically transition to any other. **3C.** Inferred rate constants (in units of 1/sec) for the same situation as in panel **B**i, but assuming that states can only transition to the adjacent mobility states. **3D.** Inferred rate constants for *clr4Δ* cells, in units of 1/sec, for the Swi6 transitions, assuming a three-state model as inferred by SMAUG. **3E.**Inferred rate constants, in units of 1/sec, assuming five biochemical states of Swi6 in the wild type cells. This model assumes that the *β* state corresponds to two chemical states: one with Swi6 bound to fully methylated H3K9me, and one with Swi6 bound to unmethylated H3K9me; the latter is assigned parameters based on the *clr4Δ* simulation shown in panel **B**.

We applied this chemical rate constant inference algorithm to the wild-type Swi6 data (shown in **Figure 1F**) to simulate transition rates between the different mobility states (*α, β, γ*, and *δ*). We plotted the complete posterior distributions to compare the simulation data with the transition rates obtained using SMAUG (**Supplementary Figure 3A**). The inferred chemical rate constants from ABC match the experimental results, demonstrating that our SMAUG analysis indeed captures the relevant timescales underlying Swi6 interstate transitions. By formal model comparison (**see Methods**) we also found that the rates inferred via ABC show a change in Bayesian information criterion (BIC) of −5.4×10^3^ relative to the SMAUG rates. As a lower BIC indicates a preferred model, we see that a detailed consideration of the chemical kinetics underlying the observed transitions yields more accurate information on the transition rates alone. Similarly, we tested whether the observed transition data are better explained by a dense transition model (**Figure 3B**), in which each state undergoes direct transitions to every other state, or by a sparse model (**Figure 3C**), in which molecules can only exchange between adjacent mobility states. We observed a change in BIC of −1.8×10^4^ for the dense model relative to the sparse model, suggesting that direct transitions between all states are necessary to recapitulate our experimental observations despite these transitions being rare in some cases.

We further examined the effects of eliminating H3K9me (via *clr4* deletion) on the interstate transitions. Based on the experimental data from **Figure 2B**, we found that the transition rates between the *γ* and *δ* states do not appreciably change in *clr4Δ* cells relative to wild-type cells (**Figure 3D**). However, the transitions involving the *β* state are significantly altered, with far more rapid transitions from the *β* to the *γ* state, presumably due to a more rapid dissociation of the Swi6 chromatin-bound *β* state in *clr4Δ* (**Figure 3D**). As discussed above, our singlemolecule tracking experiments led us to infer that the *β* state consists of at least two substates corresponding to Swi6 subpopulations interacting with H3K9me0 and H3K9me1/2/3 chromatin, respectively. Through kinetic modeling, we now find that the *β* state in *clr4Δ* cells selectively captures the behavior of the H3K9me0-bound Swi6 subpopulation.

Using our fine-grained kinetic modeling approach, we calculated the properties of the experimentally undetectable H3K9me1/2/3 bound *β* state by combining the experimental results for wild-type cells with the rate constants derived from *clr4Δ* cells. Denoting the proposed substates of *β* as *β*_0_ and *β*_me_, we repeated our ABC inference on the wild-type Swi6 data in a five-state model of the system. We assumed the observed *β* state combines the *β*_0_ and *β*_me_ substates, and we constrained parameters involving transitions amongst the *β*_0_, *γ*, and *δ* states to match those for the *clr4Δ* cells. The resulting rate constants agree substantially better with the data than did the original four-state model (ΔBIC = −5.6×10^4^), yielding the final transition rates shown in (**Figure 3E**). Hence, our hypothesis of two distinct *β* states for the wild-type Swi6 data is consistent with our experimental observations and explains the data substantially better than the four-state model. In this analysis, the *β*_0_ state is an unstable intermediate, with high rates of *β*_0_➔*γ* transitions (H3K9me0 to nucleic acid binding) and *β*_0_➔*β*_me_ transitions (H3K9me0 to H3K9me binding).

Based on the rate constants of transitions between *β*_0_ and *β*_me_ and the knowledge that approximately 2% of the *S. pombe* genome consists of H3K9me nucleosomes (http://www.pombase.org), we calculated the equilibrium constant for Swi6 binding to methylated versus unmethylated histones (ratio of rate constants between *β*_0_ and *β*_me_) to be approximately 47-fold. When the oligomerized *α* state is also accounted for, the effective equilibrium constant of Swi6 for H3K9me0 versus H3K9me1/2/3 chromatin is approximately 105-fold. Hence, oligomerization substantially enhances the specificity of Swi6 binding to sites of H3K9me in the genome.

### Oligomerization directly competes with nucleic acid binding to promote Swi6 localization at sites of heterochromatin formation

Our Swi6 single molecule tracking and simulations reveal that an intricate balance of molecular engagements involving the Swi6 CD, hinge and CSD domains facilitate Swi6 localization at sites of H3K9me *in vivo*. We initially speculated that the intermediate *β* and *γ* states might accelerate transitions between the endpoint states (*α* or *δ*) by providing a favorable set of intermediates. However, through stochastic simulations, we found that there is no significant increase in mean first passage times between *α* and *δ* state, even in the absence of both intermediates (**Supplementary Table S1**). Therefore, we hypothesized that the intermediate binding states titrate Swi6 away from sites of H3K9me and provides a tunable mechanism for highly specific H3K9me localization while preserving rapid turnover. Given a cell that is replete with DNA and RNA, how does Swi6 localize in a matter of seconds at sites of H3K9me, and maintain a stable population at silenced loci even if individual molecules of Swi6 exchange rapidly? One possibility is that Swi6 CSD mediated oligomerization and phase separation promotes Swi6 recruitment at sites of H3K9me through self-association similar to how transcription associated condensates amplify protein recruitment (Wei et al., 2020). In the case of Swi6, such a mechanism would be independent of CD mediated H3K9me recognition. A second possibility is that Swi6 CD domains are both necessary and sufficient for H3K9me recognition and stable binding. Oligomerization merely enables Swi6 to explore higher-order configurations beyond dimerization. According to the second model, Swi6 mediated H3K9me recognition in cells is independent of phase separation but depends on multivalent H3K9me binding (Chong et al., 2018).

To differentiate between these two models, we engineered strains that express combinations of Swi6 wild-type and Swi6 mutant proteins (**Figure 4A**). We took advantage of the Swi6 CD^mut^ protein which has an inactive CD domain but retains an intact CSD oligomerization domain. We co-expressed mNeonGreen-Swi6 protein in cells that also express PAmCherry-Swi6 CD^mut^. In total, these cells express two isoforms of Swi6: a CD mutant that is not capable of engaging with H3K9me chromatin and a wild-type Swi6 protein with a functional CD domain that is capable of H3K9me binding. Since mNeonGreen and PA-mCherry emissions are spectrally distinct, we used our photoactivation approach to image single PAmCherry-Swi6 CD^mut^ molecules (red channel) after verifying the presence of discrete mNeonGreen-Swi6 foci at sites of constitutive heterochromatin (green channel). We previously showed that the lack of CD mediated H3K9me recognition completely eliminates the low mobility *α* state (**Supplementary Figure 2A**). In contrast, our SMAUG analysis revealed that 5% of PAmCherry-Swi6 CD^mut^ proteins now reside in the slow mobility α state in cells that co-express wild-type mNeonGreen-Swi6 (**Figure 4B**). Therefore, CSD-mediated oligomerization makes a small but finite contribution to restore PAmCherry-Swi6 CD^mut^ α state occupancy. To confirm that the recovery of the α slow mobility state is due to CSD-dependent Swi6 interactions, we co-expressed a Swi6 CSD mutant (PAmCherry-Swi6 CSD^mut^), which is unable to oligomerize, together with the wildtype mNeonGreen-Swi6 protein. The co-expression of wild-type Swi6 protein fails to restore any measurable occupancy of PAmCherry-Swi6 CSD^mut^ protein in the low mobility *α* state (**Figure 4C**). Suppressing nucleic acid binding results in the increased occupancy of Swi6 molecules in the *α* and *β* states (**Figure 2C**). We hypothesized that eliminating nucleic acid binding might also enhance oligomerization mediated recruitment of PAmCherry-Swi6 CD^mut^ in the low mobility *α* state. Therefore, we co-expressed PAmCherry-Swi6^hinge^ CD^mut^ proteins in cells that also express mNeonGreen-Swi6. Notably, we observed a substantial increase in the occupancy of PAmCherry-Swi6^hinge^ CD^mut^ molecules in the *α* mobility state (approximately 20%) (**Figure 4D**). Therefore, our results reveal that nucleic acid binding and Swi6 oligomerization are in direct competition. Disrupting nucleic acid binding can promote an oligomerization and CD independent mode of Swi6 localization at sites of heterochromatin formation.

**Figure 4.**
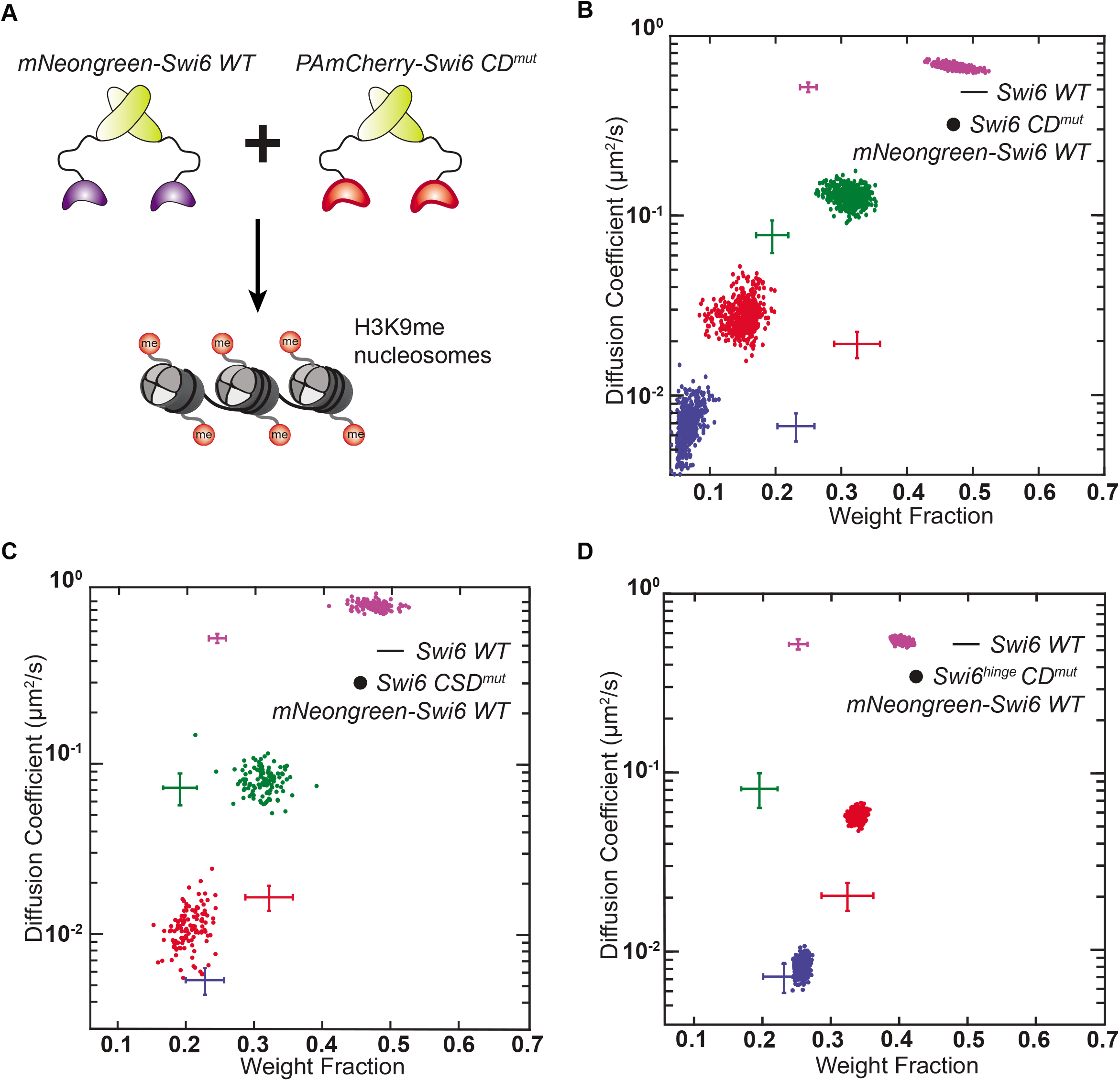
Oligomerization directly competes with nucleic acid binding to promote Swi6 localization at sites of heterochromatin formation. **4A.** Schematic representation of Swi6 co-expression experiments. The PAmCherry-Swi6 CD^mut^ and mNeonGreen-Swi6 (wild-type) proteins are expressed in the same cell and the resulting mobility states are measured using single-particle tracking of individual PAmCherry-Swi6 CD^mut^ molecules. **4B.** Average single-molecule diffusion coefficients and weight fraction estimates for PAmCherry-Swi6 CD^mut^ (*swi6 W104A* has a tryptophan to alanine substitution within the Swi6 chromodomain disrupts H3K9me recognition and binding) in cells that co-express mNeonGreen-Swi6. SMAUG identifies four distinct mobility states, (blue *α*, red *β*, green *γ*, and purple *δ*, respectively) for PAmCherry-Swi6 in CD^mut^ cells co-expressing mNeonGreen-Swi6. Each point represents the average single-molecule diffusion coefficient vs. weight fraction of PAmCherry-Swi6 molecules in each distinct mobility state at each saved iteration of the Bayesian algorithm after convergence. The dataset contains 13194 steps from 1900 trajectories. The wild-type Swi6 mobility states (**Figure 1F**) are provided as a reference (cross hairs). **4C.** Average single-molecule diffusion coefficients and weight fraction estimates for PAmCherry-Swi6 CSD^mut^ (*swi6 L315E*, a CSD domain mutation that disrupts Swi6 oligomerization) in cells that co-express mNeonGreen-Swi6. SMAUG identifies three distinct mobility states, (red *β*, green *γ*, and purple *δ*, respectively) for PAmCherry-Swi6 in CSD^mut^ cells co-expressing mNeonGreen-Swi6. Each point represents the average single-molecule diffusion coefficient vs. weight fraction of PAmCherry-Swi6 molecules in each distinct mobility state at each saved iteration of the Bayesian algorithm after convergence. The dataset contains 3200 steps from 1270 trajectories. The wild-type Swi6 mobility states (**Figure 1F**) are provided as a reference (cross hairs). **4D.** Average single-molecule diffusion coefficients and weight fraction estimates for PAmCherry Swi6^hinge^ CD^mut^ (CD^mut^ refers to a tryptophan to alanine mutation within the Swi6 chromodomain that disrupts H3K9me recognition and binding, Swi6^hinge^ is a mutant where 25 lysine and arginine residues in the hinge region are replaced with alanine) in cells that co-express mNeonGreen-Swi6. SMAUG identifies three distinct mobility states, (blue *α*, red *β* and purple *δ*, respectively) for PAmCherry-Swi6 in Swi6^hinge^ CD^mut^ cells co-expressing mNeonGreen-Swi6. Each point represents the average single-molecule diffusion coefficient vs. weight fraction of PAmCherry-Swi6 molecules in each mobility state at each saved iteration of the Bayesian algorithm after convergence. The dataset contains 15462 steps from 3250 trajectories. The wildtype Swi6 mobility states (**Figure 1F**) are provided as a reference (cross hairs).

### Increased chromodomain valency compensates for the disruption of Swi6 oligomerization

To uncouple the role of CSD mediated oligomerization from H3K9me recognition, we replaced the Swi6 CSD oligomerization domain with a glutathione-S transferase (GST) tag (**Supplementary Figure 4A**). Overall, the GST fusion construct is expected to maintain Swi6 dimerization while eliminating higher-order CSD mediated oligomerization and phase separation (Tatavosian et al., 2019). Therefore, the GST fusion construct allows us to test whether phase separation is necessary or if weak H3K9me CD recognition is sufficient for proper Swi6 localization in living cells. We refer to this hybrid protein construct as PAmCherry-Swi6^1XCD-GST^ since the newly engineered protein has only one intact CD. We expressed PAmCherry-Swi6^1XCD-GST^ in *S. pombe* cells and measured their dynamics. Following a strong 406-nm activation pulse, we imaged the ensemble of PAmCherry-Swi6^1XCD^-GST molecules in the *S. pombe* nucleus. We observed a diffuse distribution within the nucleus in contrast to wild-type-Swi6, which exhibits prominent clusters (**Supplementary Figure 4A**). Also, the PAmCherry-Swi6^1XCD^-GST single-molecule trajectories are best described by three mobility states (**Supplementary Figure 4B**). The slow mobility state *α* that corresponds to stable Swi6 binding is absent while molecules redistribute across the *β, γ* and *δ* mobility states. In addition, we also estimated transition probabilities for the different mobility states based on our experimental data. Much like our observations of wild-type Swi6 dynamics in *clr4Δ* cells, PAmCherry-Swi6^1XCD^-GST molecules predominantly reside in and transition between the *γ* and *δ* fast-mobility states with only rare transitions to the *β* state (**Supplementary Figure 4C**). These findings reinforce our earlier assignments of the various mobility states and show that a CD dimer alone is insufficient for stable Swi6 binding at sites of heterochromatin formation. Hence, a dimer of Swi6 molecules fails to localize at sites of H3K9me and is instead titrated away by non-specific DNA and/or chromatin dependent interactions. These observations are in contrast to previous studies of the mammalian HP1 isoform, HP1β, in which case replacing the CSD domain with GST had no impact on HP1β localization (Hiragami-Hamada et al., 2016). However, unlike Swi6, HP1β exhibits a reduced oligomerization capacity and fails to form condensates *in vitro* with DNA (Hiragami-Hamada et al., 2016; Keenen et al., 2020).

Next, we added a second CD domain to the existing PAmCherry-Swi6^1XCD^-GST construct to generate PAmCherry-Swi6^2XCD^-GST. The homodimerization of PAmCherry-Swi6^2XCD^-GST results in an engineered Swi6 protein that has a precise two-fold increase in CD valency relative to wild-type Swi6 and PAmCherry-Swi6^1XCD^-GST (**Figure 5A**). We used high-intensity 406-nm illumination to activate the ensemble of PAmCherry-Swi6^2XCD^-GST molecules and acquired *z*-sections at 561nm (**Figure 5C**). We observed prominent foci of PAmCherry-Swi6^2XCD^-GST molecules which qualitatively resembles wild-type Swi6 foci in cells (**Figure 5B**). Comparing the distribution of foci numbers per cell reveals a skew in the distribution with a more significant proportion of cells that exhibit three to five foci in the case of wild-type Swi6 compared to the 2XCD-GST fusion construct (**Figure 5D**).

**Figure 5.**
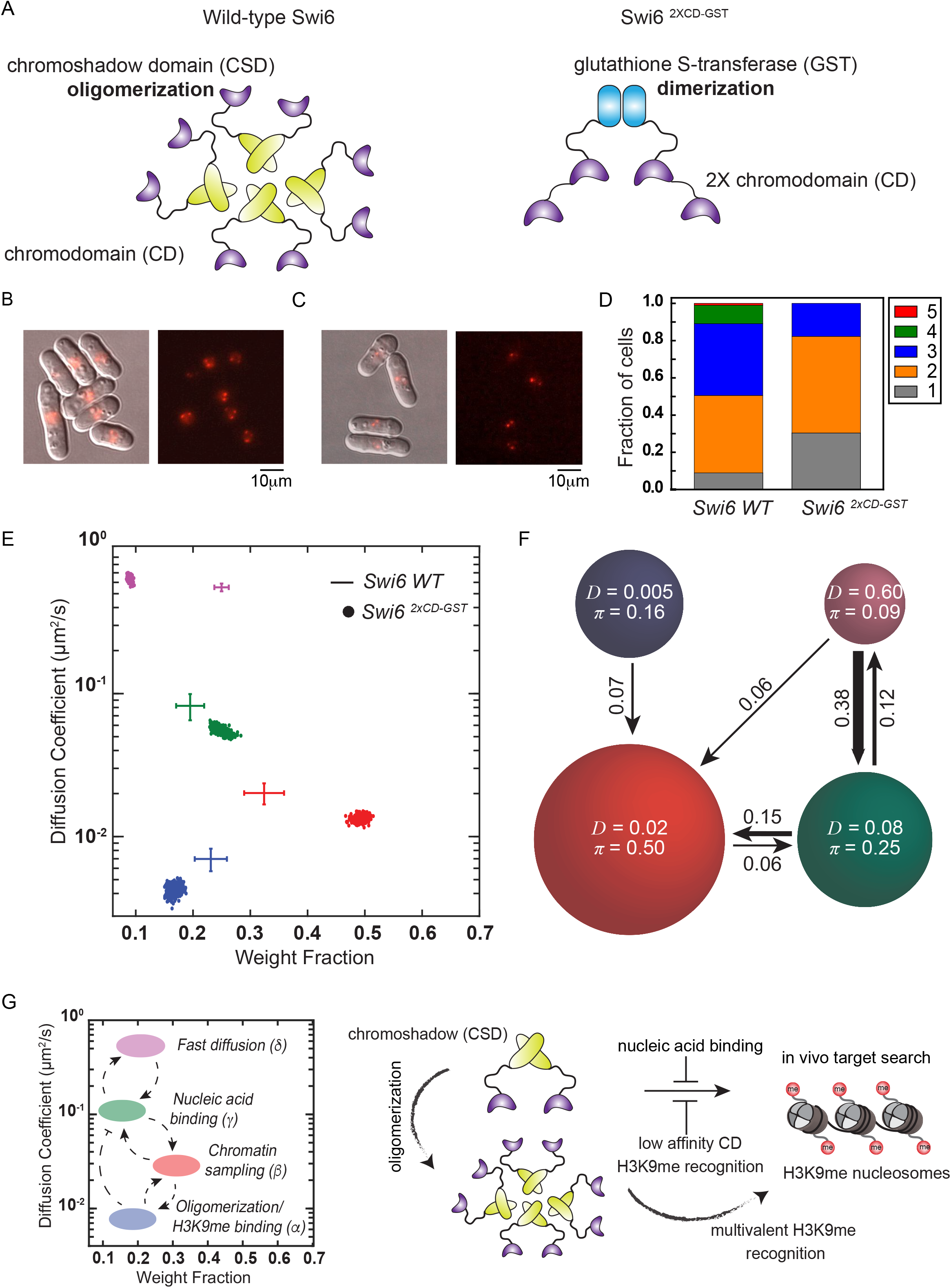
Tandem chromodomains are both necessary and sufficient for H3K9me recognition and Swi6 localization at heterochromatin. **5A.** Schematic representation of CSD mediated oligomerization of Swi6 (left) and GST mediated dimerization of an engineered Swi6^2XCD-GST^ mutant consisting of two tandem chromodomains (right). **5B.** Overlay of differential interference contrast (DIC) and epi-fluorescence images of collection of PAmCherry-Swi6 molecules that are simultaneously activated using a high 405 nm excitation power and imaged using 561 nm excitation (left panel). Epi-fluorescence image of collection of PAmCherry-Swi6 molecules simultaneously activated with high 405 nm excitation and imaged with 561 nm (right panel). The images are a maximum intensity projection of a Z-stack consisting of 13 images acquired at 250 nm z-axis intervals (scale bar 10 μm). **5C.** Overlay of differential interference contrast (DIC) and epi-fluorescence image of collection of PAmCherry-Swi6^2XCD-GST^ molecules which are simultaneously activated using a high 405 nm excitation pulse and imaged using 561 nm excitation (left panel). Epi-fluorescence image of collection of PAmCherry-Swi6^2XCD-GST^ molecules activated with high 405 nm excitation and imaged with 561 nm (right panel). The images are a maximum intensity projection of a Z-stack consisting of 13 images acquired at 250 nm z-axis intervals (scale bar 10 μm). **5D.** Distribution of the number of photoactivated PAmCherry clusters in PAmCherry-Swi6 and PAmCherry-Swi6^2XCD-GST^ expressing cells. Cells expressing wild-type Swi6 exhibit more considerable variance in the number of clusters per cell. **5E.** Average single-molecule diffusion coefficients and weight fraction estimates for cells expressing PAmCherry-Swi6^2XCD-GST^. SMAUG identifies four distinct mobility states, (blue *α*, red *β*, green *γ*, and purple *δ*, respectively) for PAmCherry-Swi6 in Swi6^2XCD-GST^ cells. Each point represents the average single-molecule diffusion coefficient vs. weight fraction of PAmCherry-Swi6 molecules in each distinct mobility state at each saved iteration of the Bayesian algorithm after convergence. The dataset contains 42382 steps from 5182 trajectories. The wild-type Swi6 clusters (**Figure 1F**) are provided for reference (cross hairs). **5F.** Based on the SMAUG identification of four distinct mobility states for PAmCherry-Swi6 in Swi6^2XCD-GST^ cells (four circles with colors as in **Figure 4E** and with average single-molecule diffusion coefficient, D, indicated in μm^2^/s), the average probabilities of transitioning between mobility states at each step are indicated as arrows between those two circles, and the circle areas are proportional to the weight fractions. Low significance transition probabilities below 0.04 are not included. The dataset contains 42382 steps from 5182 trajectories. **5G.** Model for how Swi6 locates sites of H3K9 methylation within a complex and crowded genome. Although CDs have the requisite specificity to localize at sites of H3K9me, nucleic acid binding titrates proteins away from sites of heterochromatin formation. Oligomerization stabilizes higher order configurations of the Swi6 CD domain to ensure rapid and efficient localization of Swi6 at sites of heterochromatin formation and outcompetes nucleic acid binding.

We mapped the mobility states associated with PAmCherry-Swi6^2XCD^-GST in *S. pombe*.Despite the differences in the overall number of foci, the two-fold increase in chromodomain valency completely circumvents the need for higher-order oligomerization: Swi6^2XCD^-GST fully restores the localization of Swi6 to sites of heterochromatin formation to levels that rival wildtype Swi6 (**Figure 5E**). Importantly, we measured a slow mobility state (*α* state) population of ~20%, similar to that of the wild-type Swi6 protein (**Figure 1F**). Besides, there is a substantial increase in the *β* intermediate sampling state, indicating that PAmCherry-Swi6^2XCD^-GST exhibits increased chromatin association. We confirmed that the localization of PAmCherry-Swi6^2XCD^-GST within the genome depends exclusively on H3K9me by deleting Clr4. PAmCherry-Swi6^2XCD^-GST expressing cells lacking *clr4+* exhibit a complete loss of the slow mobility α state and a concomitant decrease in the β intermediate mobility state from 50% to 12% (**Supplementary Figure 4D**). We also introduced previously characterized Swi6 CD Loop-X mutations to suppress Swi6 CD domain-dependent oligomerization, which could potentially confound our interpretations (Canzio et al., 2013). We determined that there are no quantitative differences in the mobility states associated with PAmCherry-Swi6^2XCD^-GST Loop-X and PAmCherry-Swi6^2XCD^-GST constructs without the CD Loop-X mutation (**Supplementary Figure 4E**). Hence, the recovery of the slow mobility state in the case of PAmCherry-Swi6^2XCD^-GST depends solely on CD-dependent H3K9me recognition in the absence of higher-order oligomerization. Our results reveal that four tandem CD domains are both necessary and sufficient for the stable binding of Swi6 at sites of H3K9me. In particular, since this binding occurs in the context of a chimeric protein that is not expected to undergo phase separation (Tatavosian et al., 2019), we infer that Swi6 CD domains have the requisite specificity and affinity to localize at sites of H3K9me. Hence, oligomerization is essential to promote the assembly of higher order complexes without an explicit requirement for phase separation.

Finally, we measured transition probabilities between the different mobility states in the case of PAmCherry-Swi6^2XCD^-GST (**Figure 5F**). Strikingly, we observed that molecules rarely exchange between the H3K9me dependent *α* and *β* states, unlike what we detect in the case of the oligomerization competent, wild-type Swi6 protein. The forward and reverse transition probabilities between the *α* and *β* states decrease approximately four-fold suggesting that the PAmCherry-Swi6^2XCD^-GST protein is less dynamic. These observations reveal that while multivalent CD dependent recognition is sufficient to achieve target search *in vivo*, PAmCherry-Swi6^2XCD^-GST molecules are less dynamic and exhibit fewer binding and unbinding chromatin transitions.

## DISCUSSION

Our results reveal the molecular basis for how Swi6 specifically identifies sites of H3K9me within the *S. pombe* nucleus. Unlike previous measurements of Swi6 or HP1 proteins binding to *in vitro* nucleosome substrates, our studies reveal how Swi6 binds to H3K9me nucleosomes within a native chromatin context. Remarkably, we found that Swi6 binds to H3K9me nucleosomes with 105-fold specificity, similar to *in vitro* measurements using H3K9me3 peptides as substrates (Canzio et al., 2011)Hence, Swi6 oligomerization and the presence of multivalent H3K9me chromatin domains in the nucleus allows modified histone tails to be the primary specificity determinants of Swi6 binding. We propose that Swi6 oligomerization competes with nucleic acid binding to enable low affinity H3K9me recognition chromodomains rapidly locate sites of heterochromatin formation with high specificity and selectivity (**Figure 5G**). Swi6 oligomerization stabilizes higher-order molecular configurations that promote cooperative and multivalent chromatin recognition and binding.

Unlike earlier FRAP measurements, our model-independent super-resolution assessment of Swi6 diffusion identifies four distinct mobility states (Cheutin et al., 2004; Cheutin et al., 2003; Stunnenberg et al., 2015). Using fission yeast mutants, we have validated the biochemical attributes associated with each mobility state. Our studies highlight how the high-resolution tracking of the in vivo dynamics of single molecules in cells can capture the biochemical features of proteins even in a native chromatin context. Our studies represent a vital step towards the ultimate goal of in vivo biochemistry where the on and off rates of proteins and their substrates can be reliably and directly measured in their cellular environment. Finally, we used our measured transition rates, combined with known biochemical parameters, to infer the precise chemical rate constants governing the behavior of Swi6. The primary drivers of Swi6 mobility are free diffusion, nucleic acid binding and weak and strong H3K9me-dependent interactions. The transition probabilities reveal how each mobility state functions to sequester or titrate Swi6 molecules suggesting that altering their relative occupancy ultimately affects the H3K9me bound population of the protein. Most prominently, we find that nucleic acid binding titrates Swi6 away from sites of H3K9me while neutralizing nucleic acid binding promotes stable interactions at sites of H3K9me (*α* state) and also increases the overall chromatin-bound population of the protein (*β* state). We propose that a significant function of Swi6 oligomerization is to counterbalance these inhibitory and titratable molecular interactions which would otherwise wholly suppress H3K9me localization in cells.

Preserving the same degree of nucleic acid binding but eliminating oligomerization (Swi6^1XCD-GST^) disrupts Swi6 localization to sites of H3K9me (**Supplementary Figure 5A-B**). Simply adding a second CD restores H3K9me specific localization. Although multivalency represents a longstanding principle that promotes Swi6 recruitment to sites of H3K9me, our results reveal that a defined number of tandem CD domains are both necessary and sufficient for effective H3K9me dependent localization in cells. Although recombinant Swi6 purified from *E.coli* is predominantly a dimer (~ 83%) *in vitro*, about 10% of Swi6 molecules spontaneously form tetramers (Canzio et al., 2011). Based on our results, we hypothesize that the formation of Swi6 condensates in cells increases the local concentration of Swi6 molecules in order to shift the equilibrium from Swi6 dimers to Swi6 tetramers.

In the case of Swi6 and its mammalian homolog, HP1α, oligomerization is essential to promote the formation of chromatin condensates that exhibit liquid-like properties (Larson et al., 2017; Larson and Narlikar, 2018; Sanulli et al., 2019). Our results suggest that the presence of low-affinity CD domains could be one reason why some classes of HP1 proteins are proficient in oligomerization. The ability of specific HP1 isoforms to oligomerize promotes the multivalent recognition of H3K9me nucleosomes since the dimeric state of a protein like Swi6 is insufficient for stable chromatin binding. Although we have demonstrated that low-affinity CDs can mediate Swi6 target search in vivo, our findings raise the question as to why high-affinity CD domains are not more prevalent among HP1 proteins. It is noteworthy that the engineered version of Swi6 with two CDs and no oligomerization exhibits reduced transition rates to other intermediate states from sites of H3K9me (**Figure 5F**). Therefore, these engineered proteins, once engaged in H3K9me dependent interactions, remain in these states with little to no turnover. Our attempt to simply enhance protein localization occurs at the cost of reduced Swi6 protein exchange from sites of H3K9me. Therefore, we speculate that protein recruitment which is completely CD-dependent lacks tunability. Such types of stable binding events would also effectively impede subsequent CD domain-dependent binding events which are essential for heterochromatin establishment in addition to Swi6 binding (Motamedi et al., 2008; Sadaie et al., 2008). Instead, we propose that oligomerization and protein turnover provide opportunities for regulatory inputs either via protein-protein interactions or post-translational modifications. The formation of heterochromatin condensates, in addition to serving as mechanisms that promote epigenetic silencing, could be fundamentally involved in shifting the equilibrium states of Swi6 oligomerization to promote stable H3K9me target localization in living cells (Larson and Narlikar, 2018; Strom et al., 2017).

## ACKNOWLEDGEMENTS

We are grateful to Ahmad Khalil and Ryan Baldridge for their comments on the manuscript and all members of the Biteen, Freddolino, and Ragunathan labs for their invaluable support. We thank Marc Bühler for kindly sharing the Swi6 hinge mutant plasmid and Geeta Narlikar for kindly sharing the Swi6 ARK mutant plasmid. We also thank Danesh Moazed for sharing fission yeast strains that have been used in this study. This research was funded by a National Science Foundation Understanding Rules of Life Award (1921677) to JSB, PF and the National Institute of General Medical Sciences (NIGMS) of the National Institutes of Health under award number R35GM137832 to KR. This work used the Extreme Science and Engineering Discovery Environment (XSEDE) comet resource at the San Diego Supercomputing Center through allocation TG-MCB140220 to PF (Towns et al., 2014).

## MATERIALS AND METHODS

### Plasmids

Plasmids containing Swi6 wild-type and point mutants were constructed by modified existing pFA6a N-terminal tagging plasmids. We used Gibson based gene assembly to make inframe fusion constructs of Swi6 proteins and PAmCherry. Point mutations were introduced by designing primers using guidelines described in Quick Change mutagenesis protocols. The Swi6^hinge^ mutant was amplified and subcloned into pFA6a vectors containing PAmCherry. In the case of Swi6^1XCD-GST^ and Swi6^2XCD-GST^, we obtained synthetic DNA fragments from Twist Biosciences.

### Strains

All strains were constructed using a PCR-based gene targeting approach (Bahler et al., 1998). All strains expressing PAmCherry-Swi6 or Swi6 mutants were constructed by reintroducing PCR products in swi6*Δ* strains. All strains were genotyped using colony PCR assays. Strains expressing Swi6^2XCD-GST^ or mNeonGreen-Swi6 were made by digesting the pDual vectors with Not1 restriction enzyme to facilitate their insertion at the *leu1*+ locus in *S. pombe* cells. Cells were selected based on their ability to restore leucine auxotrophy. Other deletions of heterochromatin associated factors were achieved either by PCR-based gene targeting approaches or by a cross followed by random spore analysis and PCR based screening to select for colonies that express PAmCherry-Swi6 or PAmCherry-Swi6 mutants. All strains used in this study are listed in Table S1.

### *S. pombe* live-cell imaging

Yeast strains containing a copy of PAmCherry-Swi6 or PAmCherry-Swi6 mutants under the control of the native Swi6 promoter were grown in standard complete YES media (US Biological, cat. Y2060) containing the full complement of yeast amino acids and incubated overnight at 32°C. The seed culture was diluted and incubated at 25 °C with shaking to reach an OD_600_ ~0.5. To maintain cells in an exponential phase and eliminate extranuclear vacuole formation, the culture was maintained at OD_600_ ~0.5 for 2 days with dilutions performed at ~12-hour time intervals. To prepare agarose pads for imaging, cells were pipetted onto a pad of 2% agarose prepared in YES media, with 0.1mM N-propyl gallate (Sigma, cat. P-3130) and 1% gelatin (Millipore, cat. 04055) as additives to reduce phototoxicity during imaging. *S. pombe* cells were imaged at room temperature with a 100× 1.40 NA oil-immersion objective. First, the fluorescent background was decreased by exposure to 488-nm light (Coherent Sapphire, 200 W/cm^2^ for 20 – 40 s). A 406-nm laser (Coherent Cube 405-100; 102 W/cm^2^) was used for photo-activation (200 ms activation time) and a 561-nm laser (Coherent-Sapphire 561-50; 163 W/cm^2^) was used for imaging. Images were acquired at 40-ms exposure time per frame. The fluorescence emission was filtered to eliminate the 561 nm excitation source and imaged using a 512 × 512-pixel Photometrics Evolve EMCCD camera.

For the epi-fluorescence images in Figure 4 and Supplementary Figure 4, a 405-nm LED light source (Lumencor SpectraX) at 25 mW/nm (100% power) was used to photoactivate cells and a 561 nm LED was used to image them subsequently. Images were collected with a 100-ms exposure time per frame with a 100× 1.45 NA oil-immersion objective using a Photometrics Prime95B sCMOS camera.

### Single-molecule trajectory analysis with SMAUG algorithm

Recorded Swi6-PAmCherry single-molecule positions were detected and localized with 2D Gaussian fitting with home-built MATLAB software as previously described, and connected into trajectories using the Hungarian algorithm (Munkres, 1957; Rowland and Biteen, 2017). These single-molecule trajectory datasets were analyzed by a non-parametric Bayesian framework to reveal heterogeneous dynamics (Karslake et al., 2020). This SMAUG algorithm uses non-parametric Bayesian statistics and Gibbs sampling to identify the number of distinct mobility states, *n*, in the single-molecule tracking dataset in an iterative manner. It also infers the parameter such as weight fraction, *π*_i_, and effective diffusion coefficient, *D*_i_, for each mobility state (*i*={1,2,3 *n*}), assuming a Brownian motion model. To ensure that even rare events would be captured, we collected more than 10,000 steps in our single-molecule tracking dataset for each measured strain, and we ran the algorithm over >10,000 iterations to achieve a thoroughly mixed state space. The state number and associated parameters were updated in each iteration of the SMAUG algorithm and saved after convergence. The final estimation (e.g., **Figure 1F**) shows the data after convergence for iterations with the most frequent state number. Each mobility state, *i*, is assigned a distinct color, and for each saved iteration, the value of *D_i_* is plotted against the value of *π_i_*. The distributions of estimates over the iterations give the uncertainty in the determination of *D_i_*. Furthermore, the transition probabilities (e.g., **Figure 1G**) give the average probability of transitioning between states from one step to the next in any given trajectory.

### Clustering Analysis for the Swi6 Distributions

The spatial pattern (i.e., dispersed, clustered, or homogenously distributed and at what scale) of each mobility state was investigated using Ripley’s *K* function (Ripley, 1976):

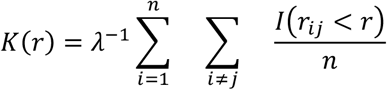

where *r* is the search radius, *n* is the number of points in the set, *λ* is the point density and, is the distance between the *i*^th^ and *j*^th^ point. *I* is an indicator function (1 when true, and 0 when false). For convenience, we further normalized K(*r*) to attain Ripley’s H function:

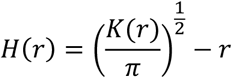

This *H*(*r*)= 0 for a random distribution, *H*(*r*)>0 for a clustered distribution pattern, and *H*(*r*) < 0 for a dispersed pattern. The maximum of H(*r*) approximately indicates the cluster size (Kiskowski et al., 2009). The cross-correlation between different states was studied with the same method. In all analysis, the nucleus was approximated as a circle to determine the area and perform edge correction (Goreaud and Pélissier, 2009). We calculated H(*r*) for each cell, then we consolidated data from different cells into an overall H(*r*) from the average across all cells weighted by the point density.

To eliminate effects from the intrinsic spatial correlation between steps that come from the same trajectories, we simulated diffusion trajectories with similar confined area size, average track length, and overall density as experimental trajectories by drawing step lengths from the step size distribution of the corresponding experiment steps. These trajectories are random in the initial position and step direction. We calculated a 4-state H(*r*) distribution for trajectories simulated corresponding to the WT Swi6 dataset, (**Supplementary Figure 1C**). To eliminate the contribution of the in-track autocorrelation of steps in H(*r*), we subtracted H(*r*) of the randomly simulated trajectories from H(*r*) of the experimental data for each mobility state. The same H(*r*) simulation and subtraction were carried out for all Ripley autocorrelation analyses.

### Fine-grained chemical rate constant inference

To calculate the chemical rate constants underlying the observed *in vivo* Swi6 transitions, we applied Bayesian inference making use of an approximated likelihood function to account for the stochasticity of the inter-state transitions occurring with the small numbers of Swi6 molecules present in living cells. This inference framework—approximate Bayesian computation (ABC)— has previously been applied to other inference problems in chemical kinetics (Beaumont et al., 2002; Csilléry et al., 2010). The overall framework for our ABC procedure is shown in Figure S6: at each iteration, rate constants are evaluated by simulating 10^5^ possible experimental transition matrices. Multinomial distributions are then fit to the probabilities of transitions (details below). Calculation of the probability that the experimentally observed state-state transition data could have occurred given these distributions yields the approximated likelihood, which is then used in standard Bayesian inference with the posterior distribution on the parameters obtained via Metropolis Markov-chain Monte Carlo. We chose an improper Jeffries prior: P(rate) ∝ 1/rate because the real chemical rate constants in this system are unknown.

Obtaining accurate likelihoods for the ABC procedure described above requires a simulation framework capable of running individual stochastic simulations of how the system might behave over a single experimental timestep given a particular set of starting populations of transition rates. We first simulate our system by dividing the experimental time (0.04 sec) into 100-time steps *δt* = 4×10^-4^sec. We confirmed that this time step was small enough to satisfy our one transition per time step assumption by computing an approximate probability that two transitions would occur to the same molecule over this time step: a molecule starting at the most mobile state had a probability of 0.58 of being in that state after 40 ms, which implies that the probability density of leaving was exponentially distributed with parameter 13.6/sec. To give an upper bound on the probability that a molecule would transition at least twice over a time step, we calculated this probability assuming all states were equally mobile, which gave us the probability of observing at least two transitions in a time step as:

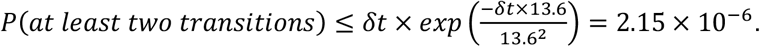

The probability of two transitions occurring in one time step even once in an experiment with 1500 molecules is therefore less than 0.3%.

At the beginning of each simulation, we assumed that the Swi6 molecules were partitioned across states according to their equilibrium populations from SMAUG. For each partition, we executed each time step based on Gillespie’s method for converting deterministic chemical rate constants to stochastic molecular behavior (Gillespie, 1977), assuming that our time step would be small enough that every particle could transition at most once. First, at each timestep, for a given diffusion state A, we calculated the probability that a given particle would transition out of that state as 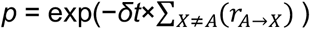, where *ρ_A→X_* is the chemical rate constant of transition from state A to state X in units of 1/sec. We sampled from a binomial distribution with transition probability *p* as given above and *n*= the number of molecules in a state to determine the number of molecules *m* which transitioned. We then determined which state each of these molecules transitioned to by generating a sample from a multinomial distribution of a size equal to *m*, where the probability that a given transitioning molecule transitions to state B is 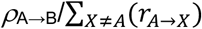. For each time step, we repeated this procedure for each Swi6 diffusion state, moving forward in time until reaching 0.04 sec, the observation time frame of diffusion experiments (schematized in **Supplementary Figure 5A**). For each calculation of the likelihood, we repeated this simulation procedure 10^5^ times, and then used the resulting matrices to fit one multinomial distribution for each initial state partition using the R package MGLM, version 0.2.0 (**Supplementary Figure 5B**). To calculate the log-likelihood of the rates, we calculated the log probability that the transition matrices observed by the experiment would have occurred given our constructed multinomial distributions and then summed them (**Supplementary Figure 5C**). Details of parameter inference for each case are given in Supplementary Text 1.To assess our models relative to each other, we calculate the Bayesian Information Criterion (BIC) (Neath and Cavanaugh, 2012; Watanabe, 2013); lower values of other information criterion indicate a preferred model. For subsequent kinetic simulation of the time required for Swi6 to transition between fully unbound and fully bound states, we calculated the mean first passage time from the *δ* to the *α* state by running our simulation 100 times with one molecule in the *δ* state and stepping in time with *δ t* = 4*10^-4^sec until our molecule ended in the *α* state and then calculating the mean of the result across the replicate simulations.

## SUPPLEMENTARY FIGURES

**Supplementary Figure 1.**
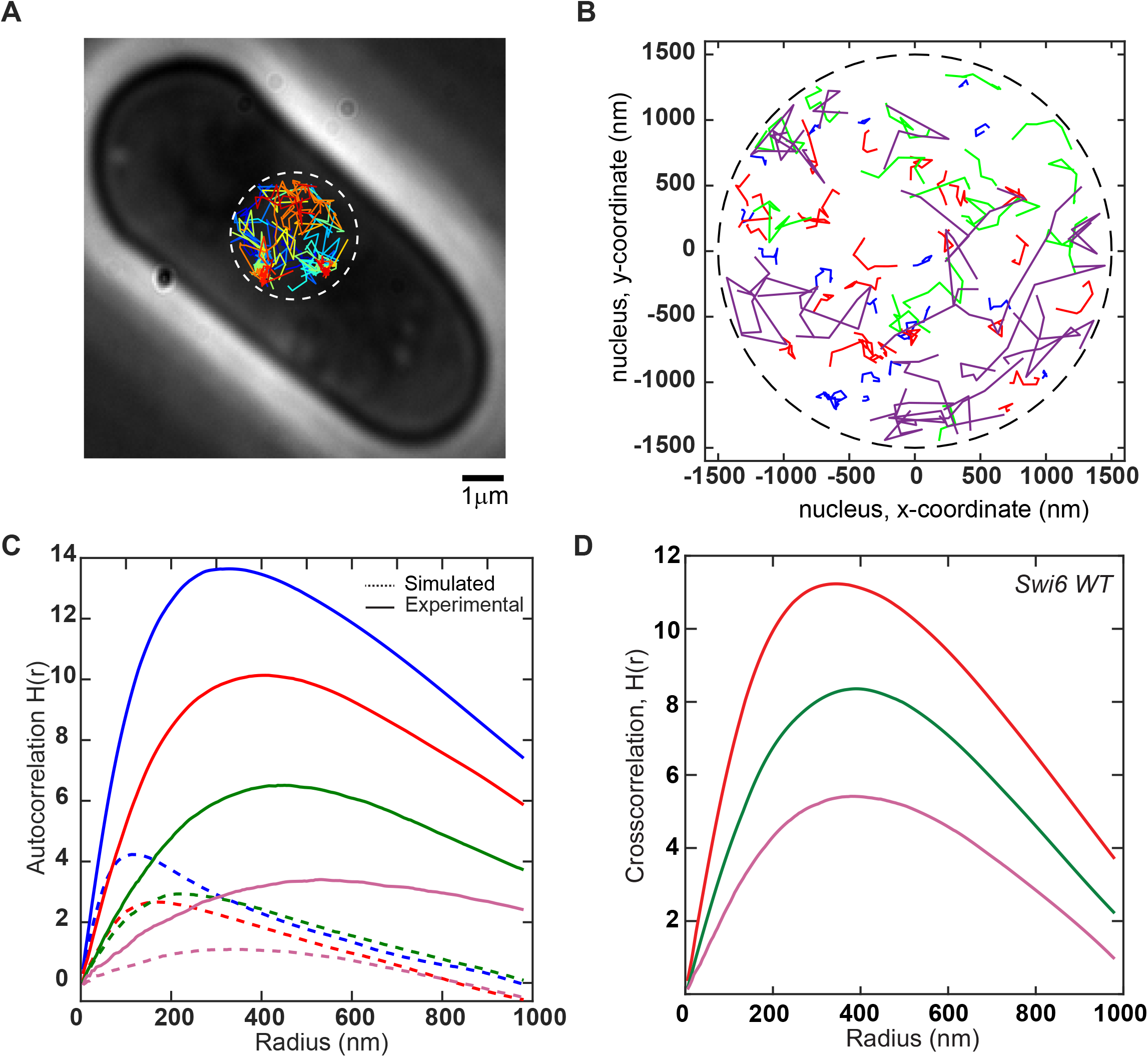
Simulations of Swi6 molecules define intervals for spatial autocorrelation analysis. **1A.** A collection of thirty-five single-particle trajectories in live *S. pombe* cells. Colors represent different single-particle trajectories that are acquired following sequential photoactivation cycles. Clusters of molecules appear at sites that correspond to constitutive heterochromatin. The heterogeneous tracks correspond to Swi6 molecules that exhibit different mobility states. **1B.** Randomly simulated trajectories for the four wild-type Swi6 mobility states. The nucleus is approximated to be a circle with radius 1.5μm as indicated by the black dashed circle. Blue, red, green and purple simulated trajectories correspond to *α, β, γ* and *δ* mobility states of wild-type Swi6. **1C.** Ripley’s H function plot for experimental trajectories and simulated trajectories of wild-type Swi6. Blue, red, green and purple lines correspond to *α*, *β*, *γ* and *δ* state. The solid line corresponds to experimentally acquired trajectories without normalization, and the dashed lines correspond to simulated trajectories shown in Figure S1B. **1D.** Ripley’s H(r) function plot for cross-correlation between *β, γ* and *δ* state in relation to the *α* state as indicated by the red, green and purple plots respectively. Simulated trajectories exhibit negligible cross-correlation values, and therefore the cross-correlation plots shown here do require any additional normalization.

**Supplementary Figure 2.**
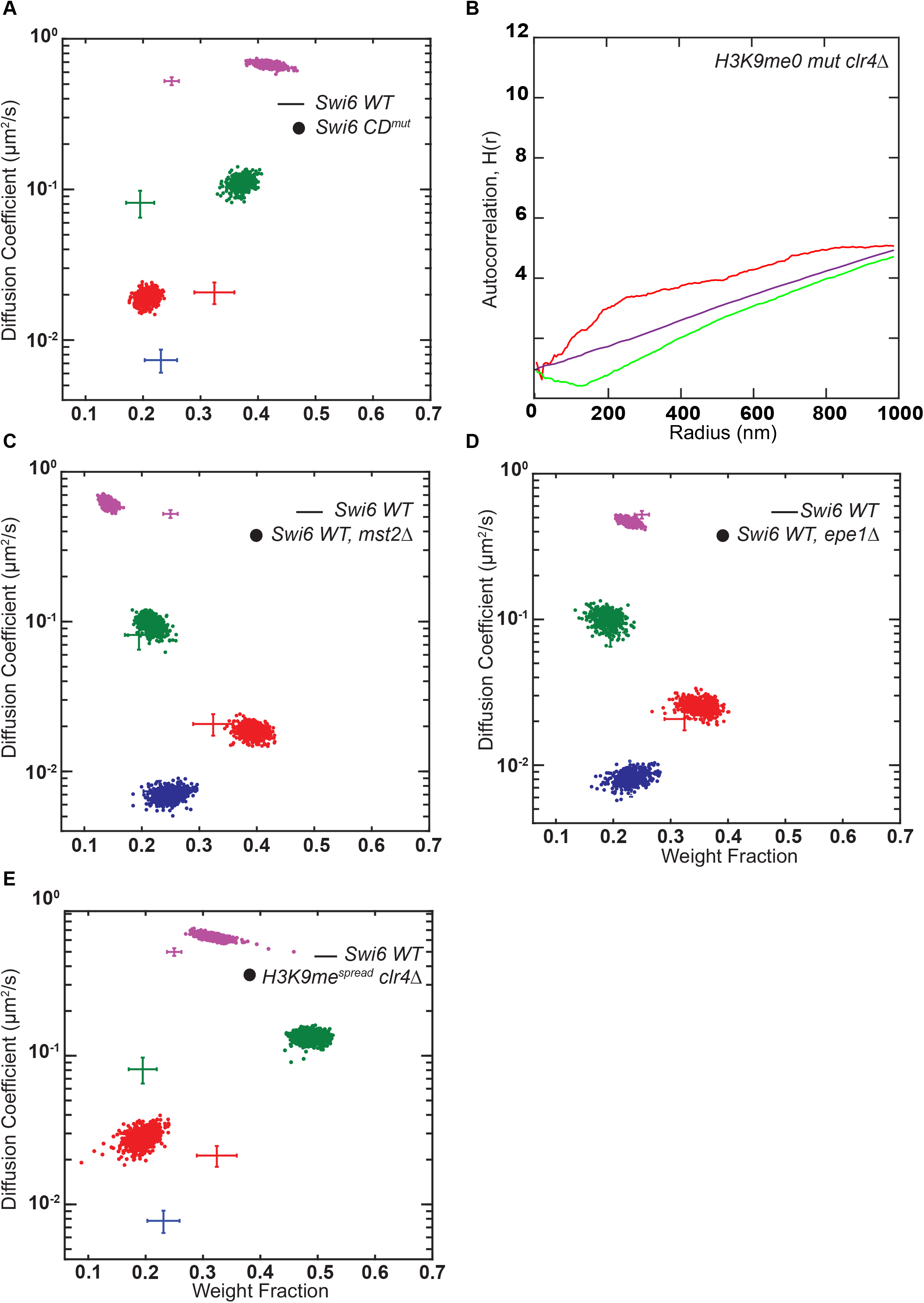
Swi6 mobility states measured in various *S. pombe* mutant backgrounds support the assignment of distinct biochemical intermediates. **2A.** Average single-molecule diffusion coefficients and weight fraction estimates for PAmCherry-Swi6 CD^mut^ expressing cells (*swi6 W104A*, which has a tryptophan to alanine mutation within the Swi6 chromodomain that disrupts H3K9me recognition and binding). SMAUG identifies three distinct mobility states, (red *β*, green *γ*, and purple *δ*, respectively) for PAmCherry-Swi6 in CD^mut^ cells. Each point represents the average single-molecule diffusion coefficient vs. weight fraction of PAmCherry-Swi6 molecules in each distinct mobility state at each saved iteration of the Bayesian algorithm after convergence. The dataset contains 10075 steps from 1624 trajectories. The wild-type Swi6 clusters (**Figure 1F**) are provided for reference (cross hairs). **2B.** The Ripley’s analysis exhibits no spatial autocorrelation for PAmCherry-Swi6 molecules in *clr4Δ* cells. H(*r*) is ≤2 for molecules in the intermediate (*β*, red, *γ*, green) and the fast (*δ*, purple) state. Each autocorrelation plot is normalized with randomly simulated trajectories from the same state (see Methods). **2C.** Average single-molecule diffusion coefficients and weight fraction estimates for PAmCherry-Swi6 molecules expressed in *mst2Δ* cells. SMAUG identifies four distinct mobility states, (blue *α*, red *β*, green *γ*, and purple *δ*, respectively) for PAmCherry-Swi6 in *mst2Δ* cells. Each point represents the average single-molecule diffusion coefficient vs. weight fraction of PAmCherry-Swi6 molecules in each distinct mobility state at each saved iteration of the Bayesian algorithm after convergence. The dataset contains 11341 steps from 1619 trajectories. The wild-type Swi6 mobility states (**Figure 1F**) are provided as a reference (cross hairs). **2D.** Average single-molecule diffusion coefficients and weight fraction estimates for PAmCherry-Swi6 molecules expressed in *epe1Δ* cells. SMAUG identifies four distinct mobility states, (blue *α*, red *β*, green *γ*, and purple *δ*, respectively) for PAmCherry-Swi6 in *epe1Δ* cells. Each point represents the average single-molecule diffusion coefficient vs. weight fraction of PAmCherry-Swi6 molecules in each distinct mobility state at each saved iteration of the Bayesian algorithm after convergence. The dataset contains 12790 steps from 2095 trajectories. The wild-type Swi6 mobility states (**Figure 1F**) are provided as a reference (cross hairs). **2E.** Average single-molecule diffusion coefficients and weight fraction estimates for PAmCherry-Swi6 molecules expressed in H3K9me*^spread^ clr4Δ (mst2Δ epe1Δ clr4Δ)* cells. SMAUG identifies three distinct mobility states, (red *β*, green *γ*, and purple *δ*, respectively) for PAmCherry-Swi6 in H3K9me*^spread^clr4Δ* cells. Each point represents the average single-molecule diffusion coefficient vs. weight fraction of PAmCherry-Swi6 molecules in each distinct mobility state at each saved iteration of the Bayesian algorithm after convergence. The dataset contains 12582 steps from 2869 trajectories. The wild-type Swi6 mobility states (**Figure 1F**) are provided as a reference (cross hairs).

**Supplementary Figure 3.**
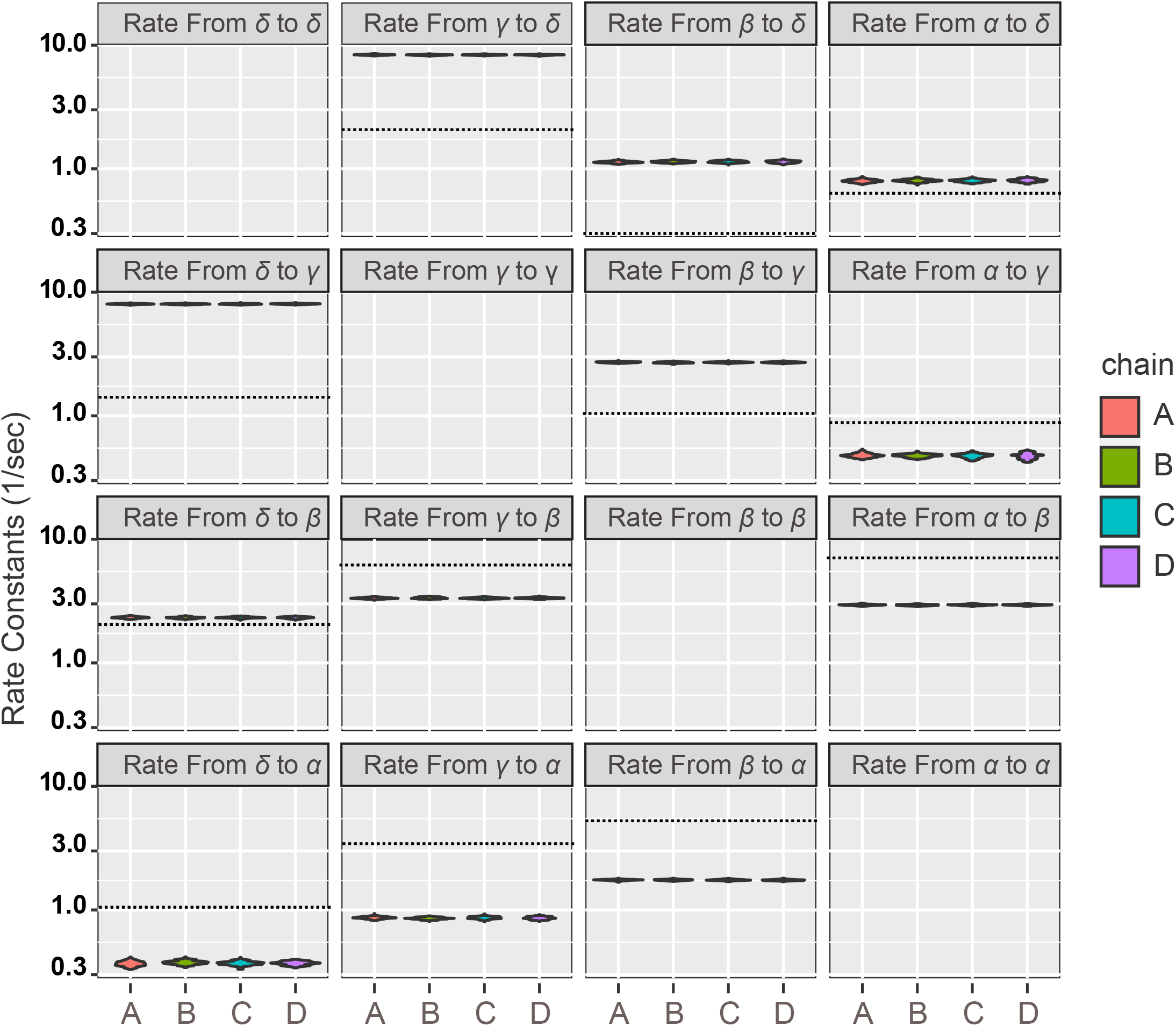
Rate Constant Model Comparisons. The posterior distributions of the Swi6 transition rate constants for the four state wild type transitions from four independent Monte Carlo chains (referred to as A-D). The dashed lines represent the rate constants that would be deduced by transforming the SMAUG transition probabilities directly into rate constants by assuming that each transition represents a single chemical transformation over the experiment.

**Supplementary Figure 4.**
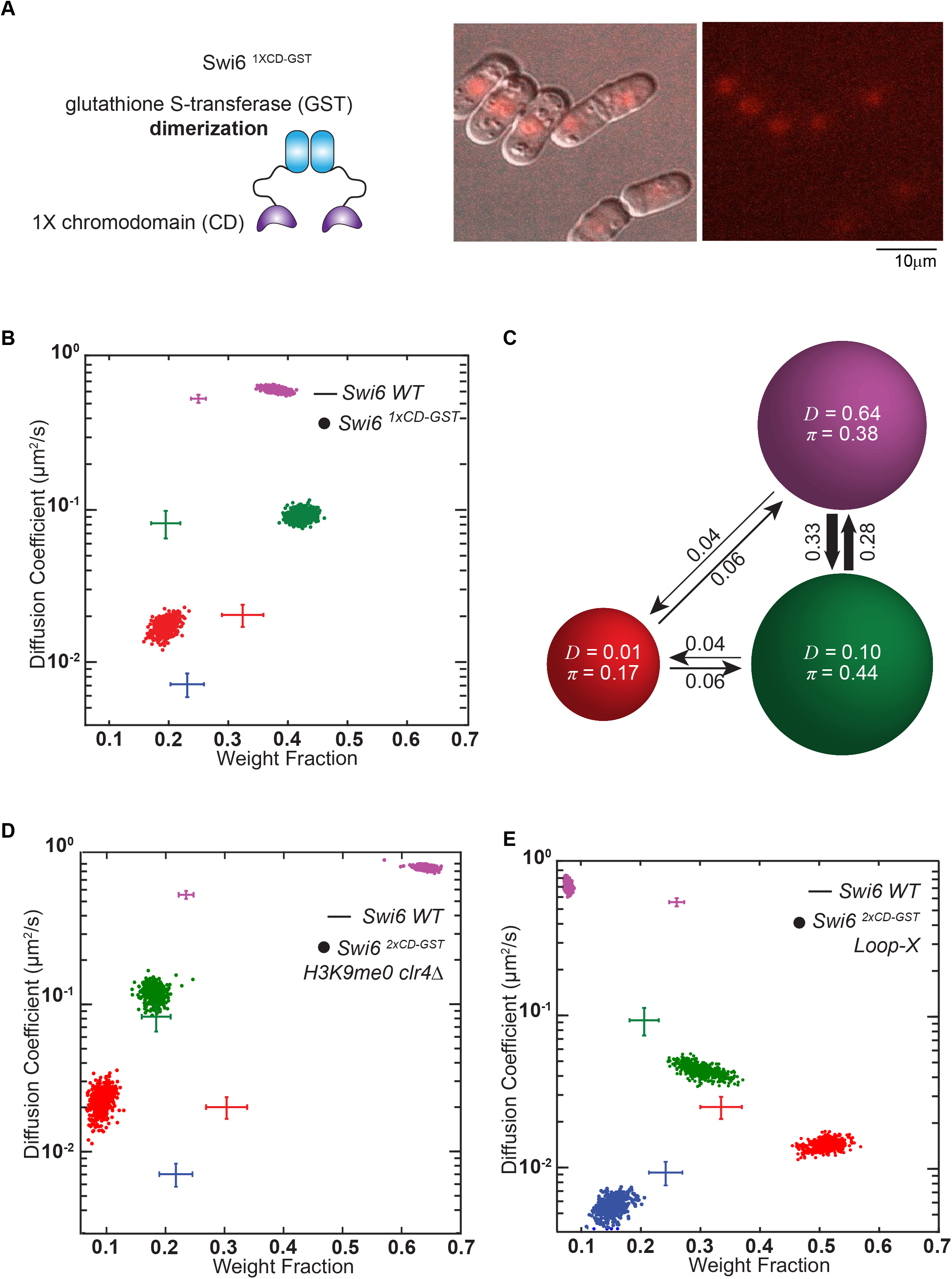
An engineered Swi6 GST fusion protein with a pair of CD domains is unable to localize at sites of H3K9me in the nucleus. **4A.** Schematic representation of an engineered, dimerization competent Swi6^1XCD-GST^ protein (left panel). Overlay of differential interference contrast (DIC) and epi-fluorescence image of collection of PAmCherry-Swi6^1XCD-GST^ molecules which are simultaneously activated using a high 405 nm excitation pulse and imaged using 561 nm excitation (left panel). Epi-fluorescence image of collection of PAmCherry-Swi6^1XCD-GST^ molecules activated with high 405 nm excitation and imaged with 561 nm (right panel). The images are a maximum intensity projection of a Z-stack consisting of 13 images acquired at 250 nm z-axis intervals (scale bar 10 μm). **4B.** Average single-molecule diffusion coefficients and weight fraction estimates for cells expressing PAmCherry-Swi6^1XCD-GST^. SMAUG identifies three distinct mobility states, (red *β*, green *γ*, and purple *δ*, respectively) for PAmCherry-Swi6 in Swi6^1XCD-GST^ cells. Each point represents the average single-molecule diffusion coefficient vs. weight fraction of PAmCherry-Swi6 molecules in each distinct mobility state at each saved iteration of the Bayesian algorithm after convergence. The dataset contains 10400 steps from 1995 trajectories. The wild-type Swi6 mobility states (**Figure 1F**) are provided as a reference (cross hairs). **4C.** Based on the SMAUG identification of three distinct mobility states for PAmCherry-Swi6 in Swi6^1XCD-GST^ cells (three circles with colors as in **Figure 5B** and with average single-molecule diffusion coefficient, D, indicated in μm^2^/s), the average probabilities of transitioning between mobility states at each step are indicated as arrows between those two circles, and the circle areas are proportional to the weight fractions, *π*. Low significance transition probabilities below 0.04 are not included. The dataset contains 10,400 steps from 1995 trajectories. **4D.** Average single-molecule diffusion coefficients and weight fraction estimates for PAmCherry-Swi6^2XCD-GST^ expressed in H3K9me0 (*clr4Δ*) cells. SMAUG identifies three distinct mobility states, (red *β*, green *γ*, and purple *δ*, respectively) for PAmCherry-Swi6 in Swi6^1XCD-GST^ cells. Each point represents the average single-molecule diffusion coefficient vs. weight fraction of PAmCherry-Swi6 molecules in each distinct mobility state at each saved iteration of the Bayesian algorithm after convergence. The dataset contains 33432 steps from 3163 trajectories. The WT clusters (**Figure 1F**) are provided for reference (cross hairs). **4E.** Average single-molecule diffusion coefficients and weight fraction estimates for cells expressing PAmCherry-Swi6^2XCD-GST^-LoopX. SMAUG identifies four distinct mobility states, (blue *α*, red *β*, green *γ*, and purple *δ*, respectively) for PAmCherry-Swi6 in Swi6^2XCD-GST^-LoopX cells. Each point represents the average single-molecule diffusion coefficient vs. weight fraction of PAmCherry-Swi6 molecules in each distinct mobility state at each saved iteration of the Bayesian algorithm after convergence. The dataset contains 22432 steps from 3138 trajectories The WT clusters (**Figure 1F**) are provided for reference (cross hairs).

**Supplementary Figure 5.**
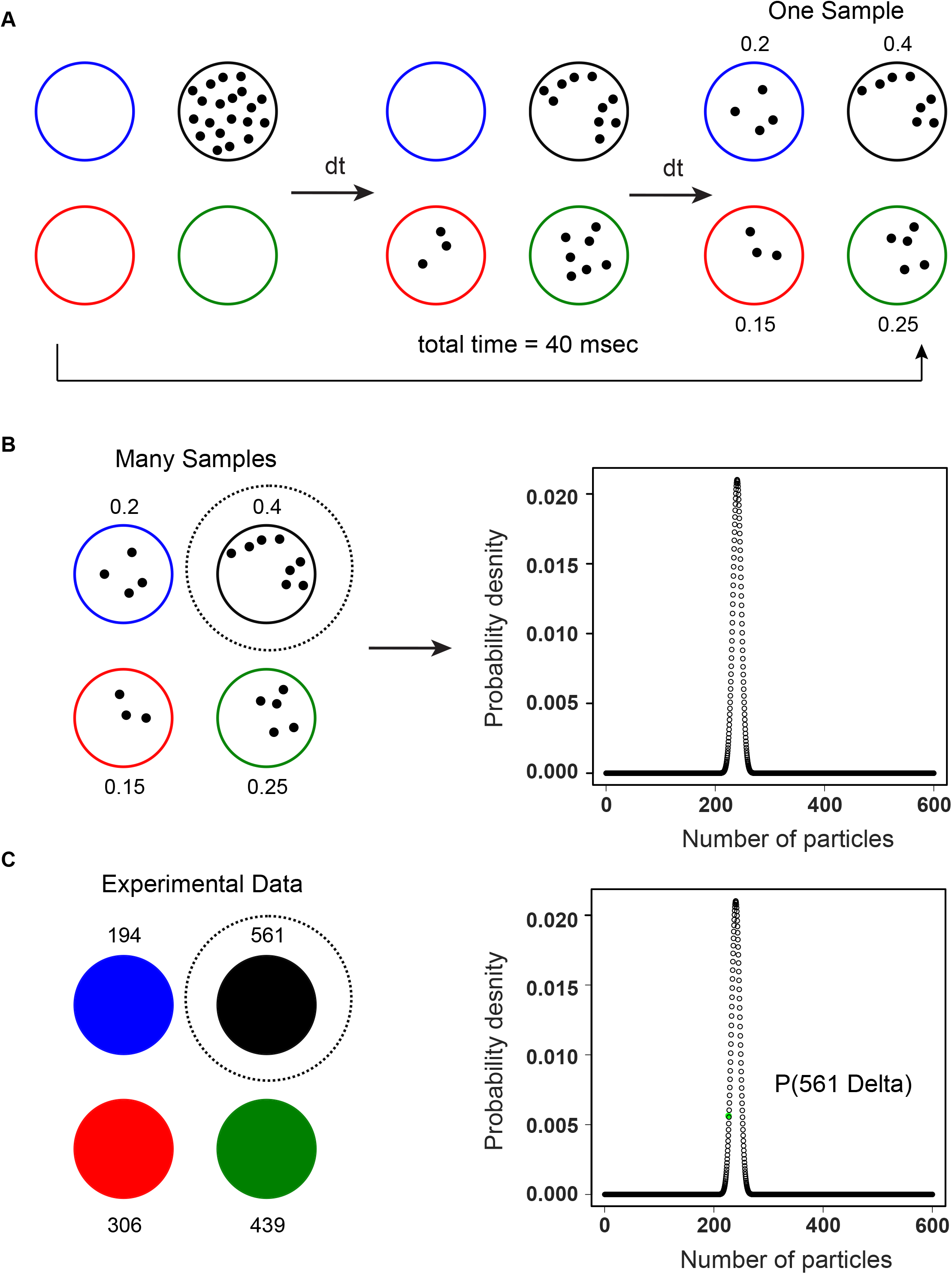
Methods for inference of fine-scale chemical rate constants. Here we schematize the steps carried out for the likelihood calculation required at each step of the Monte Carlo chain for inference of fine-grained rate parameters. **5A.** Simulation of experimental data based on fine-grained chemical rate constants. We traverse the experimental time frame of 40ms in with time step *δt* = 0.4 ms; having found that this *δt* is small enough to forbid a molecule from changing states multiple times over the course of the step (see Methods for details). At each time step, we simulate how many molecules that started in a given state-in the shown example, state *δ*-make a chemical transition over that time frame. In each simulation, this process is repeated 100 times to yield the states of the Swi6 observed at 40 ms as the experimental rate that the simulation predicts for transition rates from state *δ*. Equivalent calculations are performed for Swi6 molecules beginning in each of the other states. 10^5^ total replicate simulations (each 40 ms in duration) are performed in each case. **5B.** Construction of a likelihood function for the fates of molecules beginning in state *δ* at the end of 40 ms based on the simulations from panel **A**. A multinomial distribution is fitted to the results of the aggregated simulations from panel **A**. Similar fits are performed for each of the other beginning states to provide an empirical approximate likelihood function. **5C.** Likelihood calculation. For each starting state, we evaluate the number of experimental frames in which molecules were observed to transition to each other state after 40 ms, sum them, and then check the probability that that number would occur in the multinomial distribution constructed in panel **B**. These calculations provide likelihoods of the observed data given a particular parameter set considering each origin state separately, but in the real inference, are calculated using one multinomial distribution, as the data are not independent.

**Supplementary Table 1:** Calculated mean first passage times, in units of seconds, for one Swi6 molecule in the *δ* state to transition to the *α* state and vice versa. For the dense models where we either inferred the rates by our algorithm or where we calculated naïve rates, we also calculated the MFPT times if we omitted the *β* state, the *γ* state, or both.

